## Supporting Information for "Single Molecule Imaging Simulations with Advanced Fluorophore Photophysics"

*Dominique Bourgeois*

*Institut de Biologie Structurale, Université Grenoble Alpes, CNRS, CEA, IBS, 38044 Grenoble, France*

### Supplementary Text 1 (Application 1, Fig. 1)

#### Influence of pH

The reported pKa's of mEos4b in the green and red state are 5.5 and 5.8, respectively. Thus, at pH 8.5 the photoconversion efficiency is decreased and the red-state brightness is increased, while the opposite is true at pH 6.5. SMIS simulations suggest that at 3.5 kW/cm² better NPC images can be obtained at acidic pH. This is explained by the fact that at such readout power density the photon budget per localization is not decreased at moderately low pH (Supplementary Fig. S4C) because the median on-times are still significantly shorter than the frame time (Supplementary Fig. S4F). Strikingly, at 0.5 kW/cm² a reverse effect is observed: the image quality is better at pH 8.5. The explanation is that at pH 6.5 substantially more spots are missed (false negatives) during localization, and overall the degraded localization precision and density cannot be compensated for by the higher photoconversion efficiency.

#### Fermi profiles

Neglecting green-state photophysics, optimized Fermi profiles in principle allow maintaining a constant density of newly photoconverted red molecules along data collection. Their use permits to reduce the localization density (and therefore the numbers of spot overlaps) while maintaining the data collection time, or to reduce the data collection time while maintaining the localization density reached at constant 405-nm power. The price to pay, however, is a change in red-state photophysics along data collection resulting from the photosensitivity of dark states to 405-nm light[1]. Supplementary Fig. S5 shows that green-state photophysics compromise the possibility to maintain constant photoconversion when Fermi profiles are used, the effect being much more pronounced at 3.5 kW/cm² readout power density. Moreover, Panel 8 in Fig. 1B shows that the application of a Fermi profile while maintaining the data collection time (here 1200 s) is highly detrimental to the quality of the obtained NPC images. The reason is that with such profile (Supplementary Fig. S5E) photoconversion of most green mEos4b molecules by the 405-nm laser is delayed, while photobleaching by the 561-nm readout laser takes place at a constant rate. Hence, the photoconversion efficiency decreases substantially. This effect is much less pronounced at 0.5 kW/cm², and here there is a small gain in using a Fermi profile, attributed to the improved localization sparsity. Interestingly, the best NPC image is obtained at 3.5 kW/cm² using a Fermi profile with reduced data collection time. Here indeed, with no penalty in localization density, mEos4b molecules get photoconverted earlier, reducing green-state photobleaching and increasing the overall photoconversion efficiency. However, the data highlight that in all conditions tested, Fermi profiles result in substantially broader distributions of off-times in the mEos4b red state (Supplementary Fig. S4E), possibly compromising blinking correction algorithms employed for e.g. molecular counting.

### Supplementary Text 2 (Application 2, Fig. 2)

#### Comparison of the photophysical behaviors of mEos4b and PAmCherry

Interestingly, it can be seen that, despite its green state being highly sensitive to photobleaching by 488 nm light, the performance of mEos4b as a non-priming partner is globally similar to that of PAmCherry (Fig. 2E), which does not absorb cyan light in its non-activated state. This stems from the fact that, during the priming phase, in the absence of 405 nm light, mEos4b gets quickly shelved into its long-lived dark state, which largely protects it from photobleaching (Supplementary Fig. S8). Artificially removing in SMIS the capacity of green mEos4b to reversibly photoswitch not only results in major photobleaching during the priming phase, but also significantly increases spurious readout photoconversion during that phase, resulting in very poor colocalization (Fig. 2E and 2F). The SMIS generated datasets also highlight the impact of the different brightness of the simulated fluorescent proteins in the red states (Fig. 2D, Supplementary Fig. S6 and Supplementary Fig. S9). With a reported pKa of 7.9[2], a high fraction of the Dendra2 red chromophores remain protonated and nonfluorescent, reducing the measured photon budget per localization (Supplementary Fig. S9) and producing more blurry spots than mEos4b (Fig. 2D). With its lower intrinsic brightness as compared to mEos4b, the photon budget per localization of PAmCherry was also reduced, but this was somewhat compensated by a more complete photoactivation efficiency (Fig. 2F). Finally, as a result of their significantly different effective extinction coefficients, and therefore excitation rates by the 561 nm laser, it is interesting to notice the different levels of triplet state saturation observed in red mEos4b and Dendra2. Whereas there was almost no loss in photon budget due to intersystem crossing to T1 in the case of Dendra2 (~3%), a decrease of ~9% was noticed in the case of mEos4b (Supplementary Fig. S9).

### Supplementary Text 3 (Application 3, Fig. 3)

#### Discrepancy between simulated and experimental dSTORM data

Although a clear decrease in apparent labeling efficiency is observed as the 647 nm laser power density is increased (Fig. 3E), in line with increasing saturation of the triplet state (Supplementary Fig. S11, Supplementary Fig. S12A) from which photobleaching occurs due photon reabsorption in T1, a significant decrease is also observed at the lowest power density, also seen at the ensemble level (Supplementary Fig. S12B). Also, while the number of photons per localization decreases at high power densities, as experimentally observed, it does not rise as high as expected at lower densities (Fig. 3F). Similarly, the number of localizations per fluorophore drops, as expected, at high densities but plateaus and even decreases at lower densities (Fig. 3E), and a similar trend occurs for the global photon budget per molecule (Supplementary Fig. S12D). The highest discrepancy between the simulated and experimental data concerns the on-times (Supplementary Fig. S12C) which tend to rise in SMIS simulations instead of globally decreasing in the experimental case as the 647 nm laser power density increases.

#### Discussion on the photophysical model of Cy5

To investigate the possible origin for partial disagreement between the SMIS simulations and the experimental data, we attempted to modify some of the interconversion rates and quantum yields in the Gidi et al model[3]. First, the decrease in labeling efficiency at low power observed in the SMIS data could be assigned to thermal recovery from the $SR^-$ adduct state, which reduces the duty cycle of Cy5 switching at such low power. Thus, we assumed that in the *in cellulo* environmental conditions, the thermal recovery from SR- could be slower by a factor of ~4. Then, to increase the contrast in photon budget between low and high intensities, we reasoned that the triplet state saturation could be higher than predicted by the Gidi et al model in the experimental data, for example due to a slightly underestimated pH or concentration of β-mercaptoethanol. We also assumed that the ~4-fold superior rate of adduct formation from the singlet state, as compared to the triplet state, measured by Gidi et al[3] could be significantly lower in cellular conditions. The modified interconversion rates and quantum yields (Supplementary Table S6) provided the titration data reported in Fig. 3E-F (dashed lines). A more consistent labeling efficiency at the lowest power density was retrieved (Fig. 3E), as well as a better qualitative trend in terms of localizations per fluorophore (Fig. 3E) and photon budget (Supplementary Fig. S12D). However, the number of photons per localization (Fig. 3F) and the on times (Supplementary Fig. S12C) still deviated from the experimental data.

We reasoned that if the main dark state responsible for adduct formation and photobleaching has a lifetime inferior to the shortest frametime in the titration series (which is the case for the μs-lived triplet state), a reduction of the on times as laser illumination is augmented can only be reached if T1 saturation is lower than ~50% at the highest intensity (Supplementary Note 1). Such a condition, however, would be incompatible with the high contrast observed throughout the titration series in effective labeling densities, photons per localization and localizations per fluorophore. Thus, we conclude that a long-lived (ms) dark state is likely involved in controlling Cy5 photobleaching in dSTORM experiments. In fact, hints towards the existence of such a state has been provided experimentally[4,5], and a very long-lived triplet state had to be invoked by Diekmann et al[6] to explain their data using a simple photophysical model. Yet, T1 cannot be ms-lived under dSTORM conditions. Another argument in disfavor of the μs-lived T1 being the pivotal dark state is the huge photobleaching quantum yield >> $10^{-3}$ required in SMIS to explain the experimental data. The long-lived dark state could possibly involve one of the anionic or cationic radical states, or be generated from the cis state of Cy5, but the conclusion is that, despite its sophistication, the model of Gidi et al still appears insufficient to explain the experimental dSTORM data. More complex schemes could be tested with SMIS, but the nature of the presumed long-lived dark state remains to be elucidated.

### Supplementary Text 4 (Application 5, Fig. 5)

#### SMIS SPT data with a 5 ms frametime

Reducing the total frame time to 5 ms, the expected 2-state model was now retrieved with highest probability in the absence of fast exchange (Supplementary Fig. S17B). However, in the presence of fast exchange, a wrong 3-state model was again found, this time because the fast exchange rate becomes close to the inverse of the shortened frametime. Suppressing sensitivity of vbSPT to the fast exchange process required raising the rates to values as high as 100,000 $s^{-1}$ (Supplementary Fig. S17B). Of note, with mEos3.2, at 5 ms frame time, the localization precision limits the accuracy of the slow diffusion coefficient retrieval and no improvement is obtained by either of the techniques presented above (Supplementary Fig. S18-19).

### Supplementary Table S1

*Phototransformation quantum yields and thermal rates used for mEos4b in NPC SMIS simulations (Application 1)*

| | ***Green anionic*** | ***Green neutral*** | ***Green short-lived dark*** | ***Green long-lived dark*** | ***Green bleached*** | ***Red anionic*** | ***Red neutral*** | ***Red short-lived dark*** | ***Red long-lived dark*** | ***Red bleached*** |
|---|---|---|---|---|---|---|---|---|---|---|
| ***Green anionic*** | | Rapid Exchange | $3.5\times10^{-5}$ | $5.0\times10^{-5}$ | $2.5\times10^{-6}$ | $5\times10^{-7}$ | 0 | 0 | 0 | 0 |
| ***Green neutral*** | Rapid Exchange | | 0 | 0 | $2.5\times10^{-6}$ | $1.0\times10^{-4}$ | 0 | 0 | 0 | 0 |
| ***Green short-lived dark*** | $1\times10^{-3}$ (0.1 $s^{-1}$)* | 0 | | 0 | $2.5\times10^{-6}$ | 0 | 0 | 0 | 0 | 0 |
| ***Green long-lived dark*** | $1\times10^{-2}$ (0.001 $s^{-1}$) | 0 | 0 | | $2.5\times10^{-6}$ | 0 | 0 | 0 | 0 | 0 |
| ***Red anionic*** | | | | | | | Rapid Exchange | $5.0\times10^{-6}$ | $1.5\times10^{-5}$ | $1.0\times10^{-5}$ |
| ***Red neutral*** | | | | | | Rapid Exchange | | 0 | 0 | $1.0\times10^{-5}$ |
| ***Red short-lived dark*** | | | | | | $1.0\times10^{-4}$ (20 $s^{-1}$) | 0 | | 0 | $1.0\times10^{-5}$ |
| ***Red long-lived dark*** | | | | | | $5.0\times10^{-3}$ (0.001 $s^{-1}$) | 0 | 0 | | $1.0\times10^{-5}$ |

* Value in parentheses refer to thermally induced relaxation rates

### Supplementary Table S2

*Main parameters used in SMIS simulations (Application 1)*

| ***Simulation name*** | ***Reference*** | ***3.5 kW/cm$^2$, pH 7.5*** | ***0.5 kW/cm$^2$, pH 7.5*** | ***3.5 kW/cm$^2$, pH 8.5*** | ***3.5 kW/cm$^2$, pH 6.5*** | ***0.5 kW/cm$^2$, pH 8.5*** | ***0.5 kW/cm$^2$, pH 6.5*** | ***3.5 kW/cm$^2$, long Fermi*** | ***0.5 kW/cm$^2$, long Fermi*** | ***3.5 kW/cm$^2$, short Fermi*** |
|---|---|---|---|---|---|---|---|---|---|---|
| ***Frame time*** *(ms)* | 50 | | | | | | | | | |
| ***Number of frames*** | 25000 | | | | | | | | | 8000 |
| ***561-nm laser power density*** *[kW/cm$^2$]* | 3.5 | 3.5 | 0.5 | 3.5 | 3.5 | 0.5 | 0.5 | 3.5 | 0.5 | 3.5 |
| ***405-nm laser power density*** *[kW/cm$^2$]* | 0.002 | | | | | | | 100$^{£}$<br>2.7$^{*}$<br>1200$^{\$}$ | 100$^{£}$<br>6.7$^{*}$<br>1200$^{\$}$ | 100$^{£}$<br>5.3$^{*}$<br>360$^{\$}$ |
| ***Pixel size*** *[nm]* | 125 | | | | | | | | | |
| ***Emission filter band path*** *[nm]* | 620 ± 42.5 | | | | | | | | | |
| ***EMCCD gain*** | 200 | | | | | | | | | |
| ***EMCCD quantum efficiency*** | 0.9 | | | | | | | | | |
| ***EMCCD readout noise*** *[e$^-$]* | 74 | | | | | | | | | |
| ***Objective numerical aperture*** | 1.49 | | | | | | | | | |
| *mEos4b pH* | 7.5 | 7.5 | 7.5 | 8.5 | 6.5 | 8.5 | 6.5 | 7.5 | 7.5 | 7.5 |

£: Maximum power density
*: Fermi profile sharpness [s]
$: Fermi profile length [s]

### Supplementary Table S3

*Phototransformation quantum yields and thermal rates used for Dendra2 in primed photoconversion SMIS simulations (Application 2)*

| | ***Green anionic*** | ***Green neutral*** | ***Green T1*** | ***Green short-lived dark*** | ***Green long-lived dark*** | ***Green bleached*** | ***Red anionic*** | ***Red neutral*** | ***Red T1*** | ***Red short-lived dark*** | ***Red long-lived dark*** | ***Red bleached*** |
|---|---|---|---|---|---|---|---|---|---|---|---|---|
| ***Green anionic*** | | Rapid Exchange | $1.0\times10^{-3}$ | $3.5\times10^{-5}$ | $5.3\times10^{-5}$ | $1.0\times10^{-5}$ | $5\times10^{-7}$ | 0 | 0 | 0 | 0 | 0 |
| ***Green neutral*** | Rapid Exchange | | $1.0\times10^{-3}$ | 0 | 0 | $1.0\times10^{-5}$ | $1.0\times10^{-4}$ | 0 | 0 | 0 | 0 | 0 |
| ***Green T1*** | ($2500\ s^{-1}$) | 0 | | 0 | 0 | 0 | $1.0\times10^{-3}$ | 0 | 0 | 0 | 0 | 0 |
| ***Green short-lived dark*** | $1\times10^{-3}$ ($0.1\ s^{-1}$) | 0 | | | 0 | $1.0\times10^{-5}$ | 0 | 0 | 0 | 0 | 0 | 0 |
| ***Green long-lived dark*** | $1\times10^{-1}$ ($0.001\ s^{-1}$) | 0 | | 0 | | $1.0\times10^{-5}$ | 0 | 0 | 0 | 0 | 0 | 0 |
| ***Red anionic*** | | | | | | | | Rapid Exchange | $1.0\times10^{-3}$ | $5.2\times10^{-6}$ | $8.8\times10^{-6}$ | $5.0\times10^{-5}$ |
| ***Red neutral*** | | | | | | | Rapid Exchange | | $1.0\times10^{-3}$ | 0 | 0 | 0 |
| ***Red T1*** | | | | | | | ($2500\ s^{-1}$) | 0 | | 0 | 0 | 0 |
| ***Red short-lived dark*** | | | | | | | $1.0\times10^{-4}$ ($16\ s^{-1}$) | 0 | 0 | | 0 | $5.0\times10^{-5}$ |
| ***Red long-lived dark*** | | | | | | | $5.0\times10^{-3}$ ($0.001\ s^{-1}$) | 0 | 0 | 0 | | $5.0\times10^{-5}$ |

* Value in parentheses refer to thermally induced relaxation rates

### Supplementary Table S4

*Main parameters used in SMIS simulations (Application 2)*

| *Data collection phase* | Priming | UV |
|---|---|---|
| ***Frame time*** *(ms)* | 50 | |
| ***Number of frames*** | 10000 | 10000 |
| ***561-nm laser power density*** *[kW/cm²]* | 1 | 1 |
| ***405-nm laser power density*** *[kW/cm²]* | 0 | 0.005 |
| ***488-nm laser power density*** *[kW/cm²]* | 0.004 | 0 |
| ***730-nm laser power density*** *[kW/cm²]* | 0.5 | 0 |
| ***Pixel size*** *[nm]* | 100 | |
| ***Emission filter band path*** *[nm]* | 600 ± 20 | |
| ***EMCCD gain*** | 300 | |
| ***EMCCD quantum efficiency*** | 0.9 | |
| ***EMCCD readout noise*** *[$e^-$]* | 74 | |
| ***Objective numerical aperture*** | 1.49 | |
| ***pH*** | 7.5 | |

£: Maximum power density
*: Fermi profile sharpness
$: Fermi profile length

### Supplementary Table S5

*Phototransformation quantum yields and thermal rates used for Alexa647 (Application 3)*

| | *Fluorescent state* | *T1* | *Cationic radical* | *Anionic radical* | *Sulfur adduct* | *Bleached* |
|---|---|---|---|---|---|---|
| ***Fluorescent state*** | | **$7.1\times10^{-4}$** [$] | 0 | 0 | **$3.2\times10^{-6}$** | $5\times10^{-7}$ |
| ***T1*** | **($3.5\times10^{5}$ $s^{-1}$)** | | $1.0\times10^{-3}$ | (1000 $s^{-1}$) | **(350 $s^{-1}$)** | $5.0\times10^{-3}$ |
| ***Cationic radical*** | $1\times10^{-4}$+(**2300 $s^{-1}$**)* | 0 | | 0 | 0 | $1.0\times10^{-2}$ |
| ***Anionic radical*** | $1\times10^{-4}$+(**1170 $s^{-1}$**) | 0 | 0 | | 0 | $1.0\times10^{-2}$ |
| ***Sulfur adduct*** | **0.1**[£]+(**0.01 $s^{-1}$**) | | | | | $1.0\times10^{-5}$ |

*: Values in parentheses refer to thermally induced relaxation rates
$: Values in bold were extracted from Gidi et al[3]
£: Value calculated so that the product q×ε matches the action spectrum of Fig 4b in Gidi et al[3]

### Supplementary Table S6

*Phototransformation quantum yields and thermal rates used for Alexa647 (Application 3, modified parameters)*

| | *Fluorescent state* | *T1* | *Cationic radical* | *Anionic radical* | *Sulfur adduct* | *Bleached* |
|---|---|---|---|---|---|---|
| ***Fluorescent state*** | | **$7.1\times10^{-4}$** [$] | 0 | 0 | **$\underline{3.2\times10^{-7}}$** [%] | $3.5\times10^{-7}$ |
| ***T1*** [&] | **($\underline{1.8\times10^{5}}$ $s^{-1}$)** | | $1.0\times10^{-3}$ | (1000 $s^{-1}$) | **($\underline{867}$ $s^{-1}$)** | $4.0\times10^{-3}$ |
| ***Cationic radical*** [&] | $1\times10^{-4}$+(**2300 $s^{-1}$**)* | 0 | | 0 | 0 | $1.0\times10^{-4}$ |
| ***Anionic radical*** [&] | $1\times10^{-4}$+(**1170 $s^{-1}$**) | 0 | 0 | | 0 | $1.0\times10^{-2}$ |
| ***Sulfur adduct*** [&] | **0.1**[£]+($\underline{0.0025}$ $s^{-1}$) | | | | | $1.0\times10^{-5}$ |

&: Quantum yields are only indicative, as phototransformation rates are dictated by the product q×ε were ε is the associated extinction coefficient at the considered wavelength.
*: Values in parentheses refer to thermally induced relaxation rates
$: Values in bold were extracted from Gidi et al[3]
£: Value calculated so that the product q×ε matches the action spectrum of Fig 4b in Gidi et al[3]
%: Underlined values were adjusted from Gidi et al[3] to improve matching with experimental NPC dSTORM data

### Supplementary Table S7

*Extinction coefficients of Alexa647 (Application 3)*

| | *Fluorescent state* | *T1* | *Cationic radical* | *Anionic radical* | *Sulfur adduct* |
|---|---|---|---|---|---|
| $\varepsilon_{647\ nm}$ [$M^{-1}$×$cm^{-1}$] | 270000 | 71.5 | 2.75 | 0.09 | 0.0017 |
| $\varepsilon_{405\ nm}$ [$M^{-1}$×$cm^{-1}$] | 0 | 0 | 39240 | 5057 | 400 |

### Supplementary Table S8

*Bleaching brightness of Alexa647 (Application 3)*

| | *Fluorescent state* | *T1* | *Cationic radical* | *Anionic radical* | *Sulfur adduct* |
|---|---|---|---|---|---|
| q×$\varepsilon_{647\ nm}$ [$M^{-1}$×$cm^{-1}$] | 0.135 | 0.357 | 0.027 | 0.001 | 0 |
| q×$\varepsilon_{405\ nm}$ [$M^{-1}$×$cm^{-1}$] | 0 | 0 | 392 [&] | 51 [&] | 0.004 |

&: Photobleaching from the cationic and anionic radicals was deliberately set to high values to investigate the possible roles of these states in accelerated photobleaching. However, due to the very low level of 405 nm light used throughout data collection, the effective fraction of molecules that bleached from those states remained low (**Supplementary Fig. S11**)

### Supplementary Table S9

*Main parameters used in SMIS simulations (Application 3)*

| ***647-nm laser power density*** *[kW/cm²]* | **0.36** | **1.6** | **6.4** | **24** | **53** | **120** | **480** |
|---|---|---|---|---|---|---|---|
| ***Frame time*** *(ms)* | *500* | *110* | *28* | *7.5* | *3.5* | *1.5* | *0.38* |
| ***405-nm laser power density*** *[W/cm²] §* | 0.01£<br>125*<br>7500$ | 0.05£<br>27.5<br>1650$ | 0.2£<br>7*<br>420$ | 0.77£<br>1.87*<br>112.5$ | 1.65£<br>0.87*<br>52.5$ | 3.75£<br>0.37*<br>22.5$ | 15£<br>0.09*<br>5.7$ |
| ***Number of frames*** | 25000 | | | | | | |
| ***Pixel size*** *[nm]* | 125 | | | | | | |
| ***Emission filter band path*** *[nm]* | 700 ± 40 | | | | | | |
| ***EMCCD gain*** | 200 | | | | | | |
| ***EMCCD quantum efficiency*** | 0.9 | | | | | | |
| ***EMCCD readout noise*** *[$e^-$]* | 74 | | | | | | |
| ***Objective numerical aperture*** | 1.49 | | | | | | |
| ***Objective transmission efficiency [%]*** | 25.8 | | | | | | |
| ***Microscope optics transmission efficiency [%]*** | 80 | | | | | | |

§: Fermi profiles were applied for the 405 nm laser to keep the density of activated molecules approximately constant during data acquisition.
£: Maximum power density
*: Fermi profile sharpness [s]
$: Fermi profile length [s]

### Supplementary Table S10

*Phototransformation quantum yields and thermal rates used for Alexa647, CF680 and CF660C in Spectral Demixing SMIS simulations (Application 4)*

| | ***Fluorescent state*** | | ***Short-lived dark*** | ***Sulfur adduct*** | ***Bleached*** |
|---|---|---|---|---|---|
| ***Fluorescent state*** | | | $7.0\times10^{-5}$ | $1.0\times10^{-7}$ | $8.0\times10^{-7}$ |
| ***Short-lived dark*** | 0 | ($500\ s^{-1}$)* | | ($50\ s^{-1}$)* | $7.0\times10^{-5}$ |
| ***Sulfur adduct*** | 0.1 | ($0.0025\ s^{-1}$)* | 0 | | 0 |

* Value in parentheses refer to thermally induced relaxation rates

### Supplementary Table S11

*Main parameters used in SMIS simulations (Application 4)*

| ***Simulation name*** | ***3-color AF647/CF660C/CF680*** | ***2-color AF647/CF660C*** | ***2-color AF647/CF680*** | ***2-color AF647/AF700*** | ***1-color AF647,CF660C or CF680*** |
|---|---|---|---|---|---|
| ***Frame time*** *[ms]* | 20 | | | | |
| ***Number of frames*** | 10000 (25000$) | | | | |
| ***Number of frames*** | *10,000* | 10000<br>25000$ | *10,000* | | |
| ***561-nm laser power density*** *[W/cm²] £* | 2000- | | | | |
| ***405-nm laser power density*** *[W/cm²] £* | 0.3 | 0.3<br>0.01$<br>ramp 0.75-15& | 0.3 | | |
| ***Pixel size*** *[nm]* | 100 | | | | |
| ***Dichroic filter*** | CHROMA T685LPXR | | | SEMROCK FF695-Di01 | CHROMA T685LPXR |
| ***Emission filter Channel 1*** | CHROMA BPET68570 | | | CAIRN BP68740 | CHROMA BPET68570 |
| ***Emission filter Channel 2*** | None | | | | |
| ***Channel 2 chromatic distortion*** | [0.5, 0.7, 0.3, 0.2]£ | | | [0.7, 0.9, 0.4, 0.3]£ | [0.5, 0.7, 0.3, 0.2]£ |
| ***Fluorescence background*** *[ph/µm²/W/cm²/s]* | 100 | 100<br>10#<br>500* | 100 | | |
| ***EMCCD gain*** | 200 | | | | |
| ***EMCCD quantum efficiency*** | 0.9 | | | | |
| ***EMCCD readout noise*** *[e-]* | 74 | | | | |
| ***Objective numerical aperture*** | 1.49 | | | | |

$: Low-density dataset
&: High density dataset
£: X-shift [raster], Y-shift [raster], X-stretch [%], Y-stretch [%]
#: Low-background dataset
*: High-background dataset

### Supplementary Table S12

*Phototransformation quantum yields used for mEos3.2 in SPT SMIS simulations (Application 5)*

| | *Green anionic* | *Green neutral* | *Green short-lived dark* | *Green long-lived dark* | *Green bleached* | *Red anionic* | *Red neutral* | *Red short-lived dark* | *Red long-lived dark* | *Red bleached* |
|---|---|---|---|---|---|---|---|---|---|---|
| ***Green anionic*** | | Rapid Exchange | $3.5\times10^{-5}$ | $5.0\times10^{-5}$ | $2.5\times10^{-6}$ | $5\times10^{-7}$ | 0 | 0 | 0 | 0 |
| ***Green neutral*** | Rapid Exchange | | 0 | 0 | $2.5\times10^{-6}$ | $1.0\times10^{-4}$ | 0 | 0 | 0 | 0 |
| ***Green short-lived dark*** | $1\times10^{-3}$ ($0.1\ s^{-1}$)* | 0 | | 0 | $2.5\times10^{-6}$ | 0 | 0 | 0 | 0 | 0 |
| ***Green long-lived dark*** | $1\times10^{-2}$ ($0.001\ s^{-1}$) | 0 | 0 | | $2.5\times10^{-6}$ | 0 | 0 | 0 | 0 | 0 |
| ***Red anionic*** | | | | | | | Rapid Exchange | $1.5\times10^{-5}$ | $4.5\times10^{-5}$ | $3.0\times10^{-5}$ |
| ***Red neutral*** | | | | | | Rapid Exchange | | 0 | 0 | $3.0\times10^{-5}$ |
| ***Red short-lived dark*** | | | | | | $1.0\times10^{-4}$ ($20\ s^{-1}$) | 0 | | 0 | $3.0\times10^{-5}$ |
| ***Red long-lived dark*** | | | | | | $2.0\times10^{-2}$ ($0.001\ s^{-1}$) | 0 | 0 | | $3.0\times10^{-5}$ |

* Value in parentheses refer to thermally induced relaxation rates

### Supplementary Table S13

*Phototransformation quantum yields used for PA-JF549 (Application 5)*

| | *Off* | *Fluorescent* | *Dark* | *Bleached* |
|---|---|---|---|---|
| ***Off*** [&] | | 0.02 | 0 | 0 |
| ***Fluorescent*** | 0 | | $4.5\times10^{-6}$ | $3.0\times10^{-6}$ |
| ***Dark*** [&] | 0 | 0.02 ($35\ s^{-1}$)* | | $2.0\times10^{-6}$ |

&: Quantum yields are only indicative, as phototransformation rates are dictated by the product $q\times\varepsilon$ were $\varepsilon$ is the associated extinction coefficient at the considered wavelength.
*: Values in parentheses refer to thermally induced relaxation rates

### Supplementary Table S14

*Main parameters used in SMIS simulations (Application 5)*

| ***Simulation name*** | ***mEos3.2 -488 nm*** | ***mEos3.2 +488 nm*** | ***PA-JF549*** | ***mEos3.2 -488 nm*** | ***mEos3.2 +488 nm*** | ***PA-JF549*** |
|---|---|---|---|---|---|---|
| ***Frame time** (ms)* & | 5* (10)$ | | | 1* (4)$ | | |
| ***Number of frames*** | 10000 | | | | | |
| ***561-nm laser power density** [W/cm²]* £ | 750 | | - | 3750 | | - |
| ***555-nm laser power density** [W/cm²]* £ | - | | 750 | - | | 3750 |
| ***405-nm laser power density** [W/cm²]* £ | 2.1 | | 0.04 | 5.25 | | 0.1 |
| ***488-nm laser power density** [W/cm²]* £ | 0 | 10 | - | 0 | 25 | - |
| ***Pixel size** [nm]* | 125 | | | | | |
| ***Emission filter band path** [nm]* | 650 ± 85 | | | | | |
| ***EMCCD gain*** | 300 | | | | | |
| ***EMCCD quantum efficiency*** | 0.9 | | | | | |
| ***EMCCD readout noise** [$e^-$]* | 74 | | | | | |
| ***Objective numerical aperture*** | 1.49 | | | | | |
| ***Objective depth of focus [nm]*** | 500 | | | | | |

&: Stroboscopic mode
£: Maximum power density, flat beam profile
*: Duration of CCD/561-nm or 555-nm laser exposure
$: Duration of dark time between frames (405-nm and/or 488-nm laser exposure)

### Supplementary Table S15

*Available SMLM simulators*

| | Main purpose | Fluorophore description: Photophysical model | Fluorophore description: Spectra | SptPALM | 3D | Multicolor | PSF | Lasers | Strength | Platform |
|---|---|---|---|---|---|---|---|---|---|---|
| ***SuReSim (ref [7])*** | SMLM design and validation | 3 state | No | No | Yes | No | Scalar, 2D Gaussian, 3D Astigmatism | No | Speed; Advanced labeling | Java |
| ***SMeagol (ref [8])*** | Single particle tracking | Various predefined models | No | Yes | Yes | No | Scalar, 2D Gibson-Lanni | No | Advanced diffusion models | Matlab |
| ***TestSTORM (ref [9])*** | SMLM design and validation | 3-state | Emission wavelength | No | Yes | 2 color | Scalar/ Vectorial; 2D, 3D Astigmatism | No | Versatility, 2 color with crosstalk, polarization sensitivity | Matlab |
| ***(SR FightClub) (ref [10])*** | Input for testing localization software | 4-state | No | No | Yes | No | Experimental 2D or 3D Astigmatism, Bi-plane, Double helix | No | Realistic spot shape, spot density and signal-to-noise ratio | Java |
| ***FluoSim (ref [11])*** | Simulating protein dynamics | 2-state | No | Yes | No | No | Scalar, 2D Gaussian | No | Various modalities: SMLM, sptPALM, FRAP, FCS | C++ |
| ***VirtualSMLM (ref [12])*** | Real-time SMLM simulations | 4-state | No | No | Yes | No | Experimental 2D or 3D Astigmatism, Bi-plane, Double helix | Yes | Speed; Imaging parameters can be changed on the fly | Java |
| ***sptPALMsim (ref [13] )*** | Single particle tracking | 3-state | No | Yes | Yes | No | 3D, pupil function model | No | Multiple diffusing states | Python |
| ***SMIS (This work)*** | Advanced fluorophore photophysics | N-state | Full spectra | Yes | Yes | Multicolor | Scalar, 2D Gaussian, 3D Astigmatism, | Yes | Same input parameters as on real microscope | Matlab |

**Supplementary Table S16**

*List of main SMIS features.*

### FLUOROPHORES:

- UNLIMITED NUMBER OF FLUOROPHORES (COLORS)
- NUMBER OF MOLECULES PER COLOR
- FRACTIONAL DISTRIBUTION ON VIRTUAL SAMPLE PATTERNS
- MATURATION LEVEL, LABELING EFFICIENCY
- MINIMUM DISTANCE BETWEEN FLUOROPHORES
- LINKAGE ERROR OPTION
- FLUOROGENICITY OPTION
- PH SENSITIVITY OPTION FOR FLUORESCENT STATES (PKA, HILL COEFFICIENT)
- DIPOLE ORIENTATION
    - FIXED OR TUMBLING, REORIENTATION RATE OPTION
- PHOTOPHYSICAL STATES
    - UNLIMITED NUMBER OF PHOTOPHYSICAL STATES
    - FLUORESCENCE QUANTUM YIELD FOR FLUORESCENT STATES
    - ABSORPTION, EXCITATION AND FLUORESCENCE EMISSION SPECTRA
    - THERMAL EXCHANGE RATES BETWEEN PHOTOPHYSICAL STATES
    - PHOTOTRANSFORMATION QUANTUM YIELDS BETWEEN PHOTOPHYSICAL STATES
- DIFFUSION BEHAVIOR
    - UNLIMITED NUMBER OF DIFFUSION STATES
    - DIFFUSION COEFFICIENTS
    - CONFINED DIFFUSION ON DEFINED VIRTUAL SAMPLE PATTERNS
    - DIRECTED MOTION OPTION
    - EXCHANGE RATE BETWEEN DIFFUSION STATES
- QUANTITATIVE PALM
    - DEFINED OLIGOMERIC STATE
- FLUOROPHORE PAIRING OPTION FOR COLOCALIZATION
- FRET OPTION BETWEEN PAIRS OF FLUOROPHORES

### MICROSCOPE DESCRIPTION

- MICROSCOPE OBJECTIVE AND POINT SPREAD FUNCTION
    - NUMERICAL APERTURE
    - OBJECTIVE REFRACTIVE INDEX
    - SAMPLE REFRACTIVE INDEX
    - OBJECTIVE AND MICROSCOPE TRANSMISSION EFFICIENCIES
    - 2D OPTIONS
        - GAUSSIAN PSF
    - 3D OPTIONS
        - DEPTH OF FOCUS
        - 3D SAMPLE POSITIONING
        - GAUSSIAN OR ASTIGMATIC PSF
        - TIRF OR HILO MODES
- LASERS

        - UNLIMITED NUMBER OF LASERS
        - LASER DESCRIPTION:
            - WAVELENGTH
            - PROFILE (GAUSSIAN OR FLAT)
            - *NOMINAL POWER OR POWER DENSITY AT BEAM CENTER*
            - *INTENSITY ALONG DATA ACQUISITION (CONSTANT, RAMP, FERMI PROFILE, CUSTOM PROFILE)*
            - *POLARIZATION (CIRCULAR OR LINEAR)*
            - *TIMING (ON DURING FRAMES OR BETWEEN FRAMES)*
            - *FRAP MODE (LOCAL ILLUMINATION)*
- ACQUISITION CHANNELS AND FILTERS
    - 2-CHANNEL OPTION
        - DUAL CAMERA OR SPLIT SINGLE CAMERA
        - CHANNEL 2 DEFOCUS AND DEFORMATION OPTIONS
    - EMISSION AND DICHROIC FILTERS
        - EXPERIMENTAL FILTERS OPTION
- EMCCD CAMERA
    - EMCCD GAIN, QUANTUM EFFICIENCY, DYNAMIC RANGE, ADU OFFSET, READOUT NOISE, CLOCK - INDUCED CHARGE, CONVERSION FACTOR ($e^-$/ph, $e^-$/ADU)

---

### *DATA COLLECTION PARAMETERS*

---

- FRAME SIZE, NUMBER OF FRAMES, PIXEL SIZE, FRAME TIME, TIME BETWEEN FRAMES
- SAMPLE DRIFT OPTION

---

### *SMIS tools*

---

- CREATE FLUOROPHORE TOOL
- CREATE VIRTUAL SAMPLE TOOL
- ANALYZE RESULTS TOOL

---

### *General SMIS parameters*

---

- PARALLEL COMPUTING OPTION
- ENSEMBLE SIMULATION OPTION
- DATA ACQUISITION MONITORING
- PSF, FLUOROPHORE SPECTRA AND LASERS MONITORING TOOLS
- SAMPLING RATE OPTIMIZATION OPTION
- SPTPALM OPTIONS (SUB-FRAME DIFFUSION, DIFFUSION SAMPLING RATE)

### Supplementary Table S17

*Examples of available fluorophores in SMIS. See also Methods section for modifying available fluorophores or defining new fluorophores.*

| | *Type* | *Photophysical model* | *Spectra* | *pH dependence* | *Comment** |
|---|---|---|---|---|---|
| ***EGFP*** | FP | 3-state | FPbase$ | No | Basic, 1 dark state |
| ***EYFP*** | FP | 3-state | FPbase | No | Basic, 1 dark state |
| ***mVenus*** | FP | 2-state | FPbase | No | Basic, no blinking |
| ***mTurquoise2*** | FP | 2-state | FPbase | No | Basic, no blinking |
| ***rsEGFP2*** | RSFP | 5-state | In-house (ref [14]) | No | Includes blinking and switching |
| ***rsEGFP2 cryo*** | RSFP | 8-state | In-house (ref [15]) | No | 2 fluorescent states with corresponding blinking and bleaching |
| ***rsTagRFP*** | RSFP | 4-state | ref [16] | No | Includes blinking and switching |
| ***Simple PCFP*** | PCFP | 6-state | Based on ref [17] | Yes | Generic PCFP without blinking |
| ***mEos4b*** | PCFP | 10-state | In-house (ref [17]) | Yes | Includes blinking, switching and photoconversion |
| ***mEos4b T1*** | PCFP | 11-state | In-house (ref [17]) | Yes | Includes triplet state in the red state in addition to mEos4b model |
| ***mEos3.2*** | PCFP | 10-state | FPbase +In-house (ref [17]) | Yes | Includes blinking, switching and photoconversion |
| ***Dendra2 T1*** | PCFP | 12-state | In-house (ref [18]) | Yes | Includes blinking, switching, photoconversion and triplet state in both colors |
| ***PAmCherry1*** | PAFP | 7-state | FPbase | Yes | Includes blinking and switching in red state |
| ***PAmCherry1 T1*** | PAFP | 8-state | FPbase | Yes | Includes triplet state, blinking and switching in red state |
| ***Alexa647 STORM*** | Cyanine dye | 4-state | SpectraViewer£ | No | Ad hoc model for STORM: includes ms-lived and long-lived dark states |
| ***Alexa700 STORM*** | Near-IR dye | 4-state | SpectraViewer | No | Ad hoc model for STORM: includes ms-lived and long-lived dark states |
| ***CF680 STORM*** | Cyanine dye | 4-state | AAT Spectrum Viewer% | No | Ad hoc model for STORM: includes ms-lived and long-lived dark states |
| ***CF660 STORM*** | Cyanine dye | 4-state | AAT Spectrum Viewer | No | Ad hoc model for STORM: includes ms-lived and long-lived dark states |
| ***Cy5 full*** | Cyanine dye | 7-state | SpectraViewer and ref [3] | No | Full Cy5 model including triplet state, cis-state, anionic and cationic radicals and sulfur adduct |

| | | | | | |
|---|---|---|---|---|---|
| ***Atto647N PBS*** | Pyronine dye | 5-state | ATTO-TEC | No | Ad-hoc model for Atto647N in PBS |
| ***Atto647N GLOX*** | Pyronine dye | 5-state | ATTO-TEC | No | Ad-hoc model for Atto647N in deoxygenated medium |
| ***Atto647N STORM*** | Pyronine dye | 5-state | ATTO-TEC | No | Ad-hoc model for Atto647N in STORM buffer |
| ***Atto647N ROXS*** | Pyronine dye | 5-state | ATTO-TEC | No | Ad-hoc model for Atto647N in ROXS buffer |
| ***Atto655*** | Oxazine dye | 2-state | ATTO-TEC | No | No photophysics |
| ***Atto633*** | Pyronine dye | 2-state | ATTO-TEC | No | No photophysics |
| ***NileRed*** | Oxazine dye | 2-state | SpectraViewer | No | Fluorogenic |
| ***PA-JF549*** | Photoactivatable Rhodamine | 4-state | SMIS based on ref [19] | No | Includes photoactivation and blinking |
| ***Bead*** | - | 2-state | SMIS synthetic | No | SMIS created bead |

*: All photophysical models include bleaching
$: FPbase: https://www.fpbase.org/
£: ThermoFisher SpectraViewer: https://www.thermofisher.com/order/fluorescence-spectraviewer
%: AAT Spectrum Viewer: https://www.aatbio.com/fluorescence-excitation-emission-spectrum-graph-viewer
#: ATTO-TEC spectra: https://www.atto-tec.com/

### Supplementary Note 1: Estimation of the dependence of on times as a function of laser power density

Here, we show that, using the Cy5 photophysical model of Gidi et al[3] with the short-lived triplet state as the main source of photobleaching , a decrease in on times as the laser power density increases is incompatible with a high contrast in the number of emitted fluorescence photons between high and low intensity lasers.

The rate of triplet state formation is given by: $k_{ST} = k_{exc}^{S} \times Q_{ISC}$, where $k_{exc}^{S}$ is the excitation rate of the singlet state and $Q_{ISC}$ is the quantum yield of intersystem crossing. The characteristic time spent in the singlet state before intersystem crossing is $\tau_S = 1/\,k_{ST}$ . The rate for combined sulfur adduct formation and photobleaching from the triplet state is given by $k_{TSAB} = k_{TSA} + k_{exc}^{T} \times Q_B$ where $k_{TSA}$ is the thermally induced rate of sulfur adduct formation, $k_{exc}^{T}$ is the excitation rate of the triplet state and $Q_B$ is the photobleaching quantum yield from that state. The number of accesses to the triplet state to achieve a transition to either the sulfur adduct or the photo bleached state can then be estimated by:

$$N_T \approx 1/\tau_T \times 1/k_{TSAB}$$

with $\tau_T$ being the triplet state lifetime. As $N_T \gg 1$ when $\tau_T$ is in the microsecond range, we can estimate the on-time in [s] as:

$$t_{ON}\ [s]\ \approx N_T \times (\tau_T + \tau_S)$$

Now let's convert the on-time in units of frametimes, keeping in mind that frame times are adjusted to obtain a constant illumination dose. At a given laser power density, the frame time $F$ can thus be written: $F\ [s] = K/k_{exc}^{S}$, where $K$ is the number of excitation of the singlet state per frame. Therefore, the on-time in units of frametimes can be expressed as:

$$t_{ON}\ [f]\ \approx k_{exc}^{S}/K \times N_T \times (\tau_T + \tau_S)$$

At low laser power density, there is little photobleaching from the triplet state and we have $k_{TSAB} \approx k_{TSA}$. Thus the on time $t_{ON}^{L}\ [f]$ reduce to:

$$t_{ON}^{L}\ [f]\ \approx k_{exc,L}^{S}/K \times 1/k_{TSA} \times \left(1 + {\tau_S^L}/{\tau_T}\right)$$

with $\tau_S^L$ being the characteristic time before switching to the triplet state and $k_{exc,L}^{S}$ the excitation rate under low intensity illumination. In contrast, at the highest laser power density, photobleaching predominates so that $k_{exc}^{T} \times Q_B = \alpha \times k_{TSA}$ with $\alpha > 1$ (in the case of Diekmann et al[6], $\alpha \approx 3$ at 480 kW/cm²). Thus the on time $t_{ON}^{H}\ [f]$ is expressed as:

$$t_{ON}^{H}\ [f]\ \approx k_{exc,H}^{S}/K \times 1/(\alpha + 1)k_{TSA} \times \left(1 + {\tau_S^H}/{\tau_T}\right)$$

Now the ratio of the on times at low and high laser eliminations is given by:

$$r_{L/H} = \frac{t_{ON}^{L}\,[f]}{t_{ON}^{H}\,[f]} \approx (\alpha+1)\times\frac{k_{exc,L}^{S}\times\left(1+{\tau_S^L}/{\tau_T}\right)}{k_{exc,H}^{S}\times\left(1+{\tau_S^H}/{\tau_T}\right)} = (\alpha+1)\times\frac{1+{\tau_T}/{\tau_S^L}}{1+{\tau_T}/{\tau_S^H}}$$

At low intensity, we have ${\tau_T}/{\tau_S^L} \ll 1$, as there is low triplet state saturation. Thus, the expression above simplifies to:

$$r_{L/H} = (\alpha+1)\times\frac{\tau_S^H}{\tau_S^H+\tau_T}$$

For $r_{L/H} > 2$ and $\alpha \approx 3$, as observed experimentally, we arrive at the following constraint:

$$\tau_S^H > \tau_T$$

This condition implies a rather low triplet state saturation of < 50% of the highest illumination power density, which is incompatible with the high contrast in photon detection between low and high laser intensities.

### Supplementary Figures

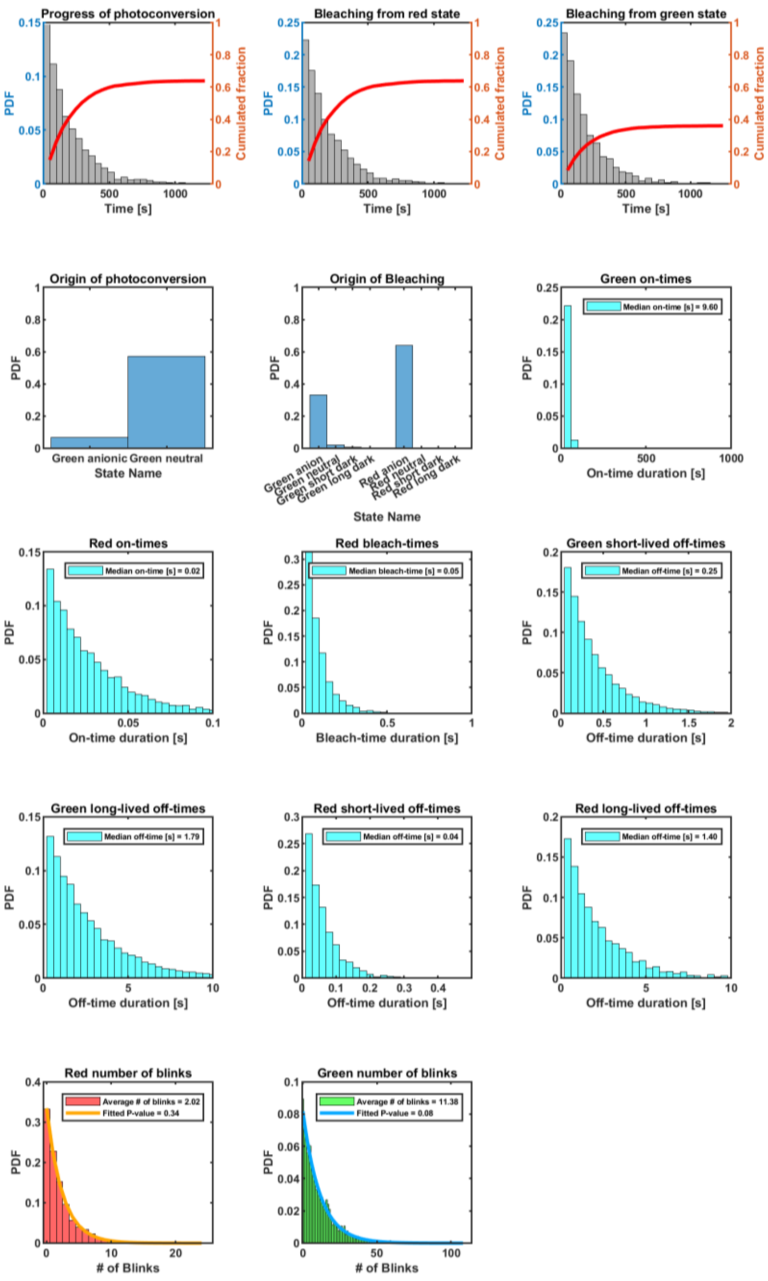


### Supplementary Fig. S1

Ground-truth photophysical behavior of mEos4b in SMIS simulations employing the model of Fig. 1A. The case of the simulation in Fig. 1C, Panel 2 is shown (3.5 kW/cm$^2$, pH 7.5). PDF: probability density function. P-value: probability of bleaching normalized by the probability of transitioning to either bleached or blinked states.

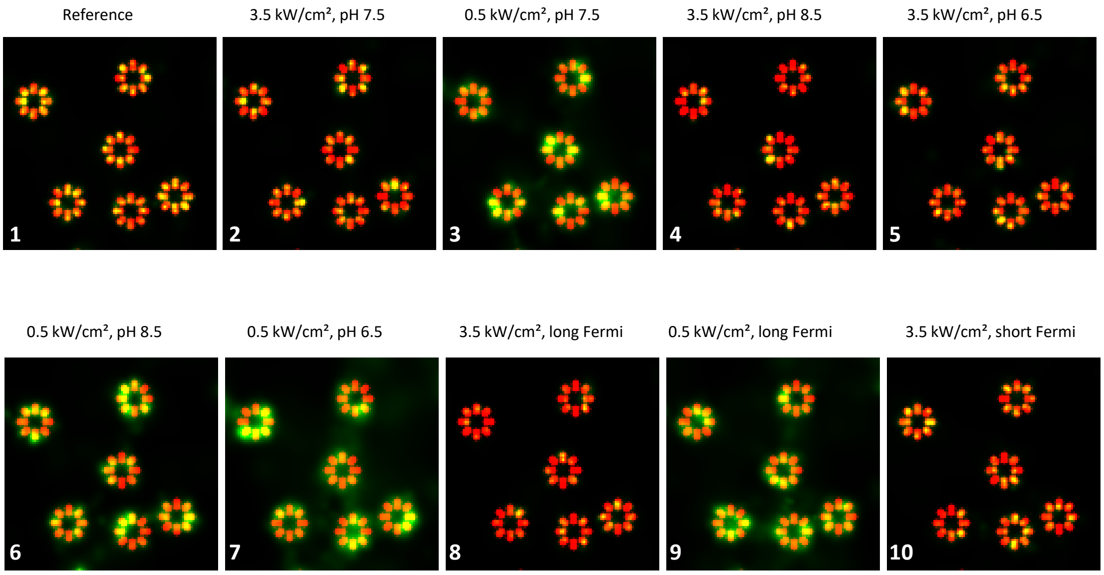


### Supplementary Fig. S2

Blend images obtained by combining rendered NPC images shown in Fig. 1C and a ground truth NPC image assuming 100% labeling efficiency and Nup96 Gaussian spots of ~10 nm fullwidth at half maximum (FWHM). The panel numbers shown correspond to those in Fig. 1C. Red color is a marker for incomplete labeling due to lack of mEos4b photoconversion or detection. Green color is a marker for degraded localization precision.

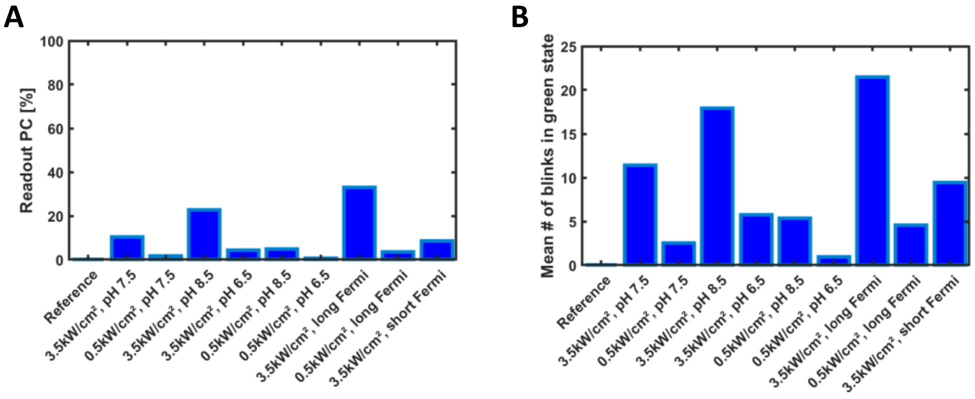


### Supplementary Fig. S3

Percentage of mEos4b readout photoconversion (A) and mean number of blinks before photoconversion (B) in the performed SMIS simulations. Values extracted from ground truth SMIS data.

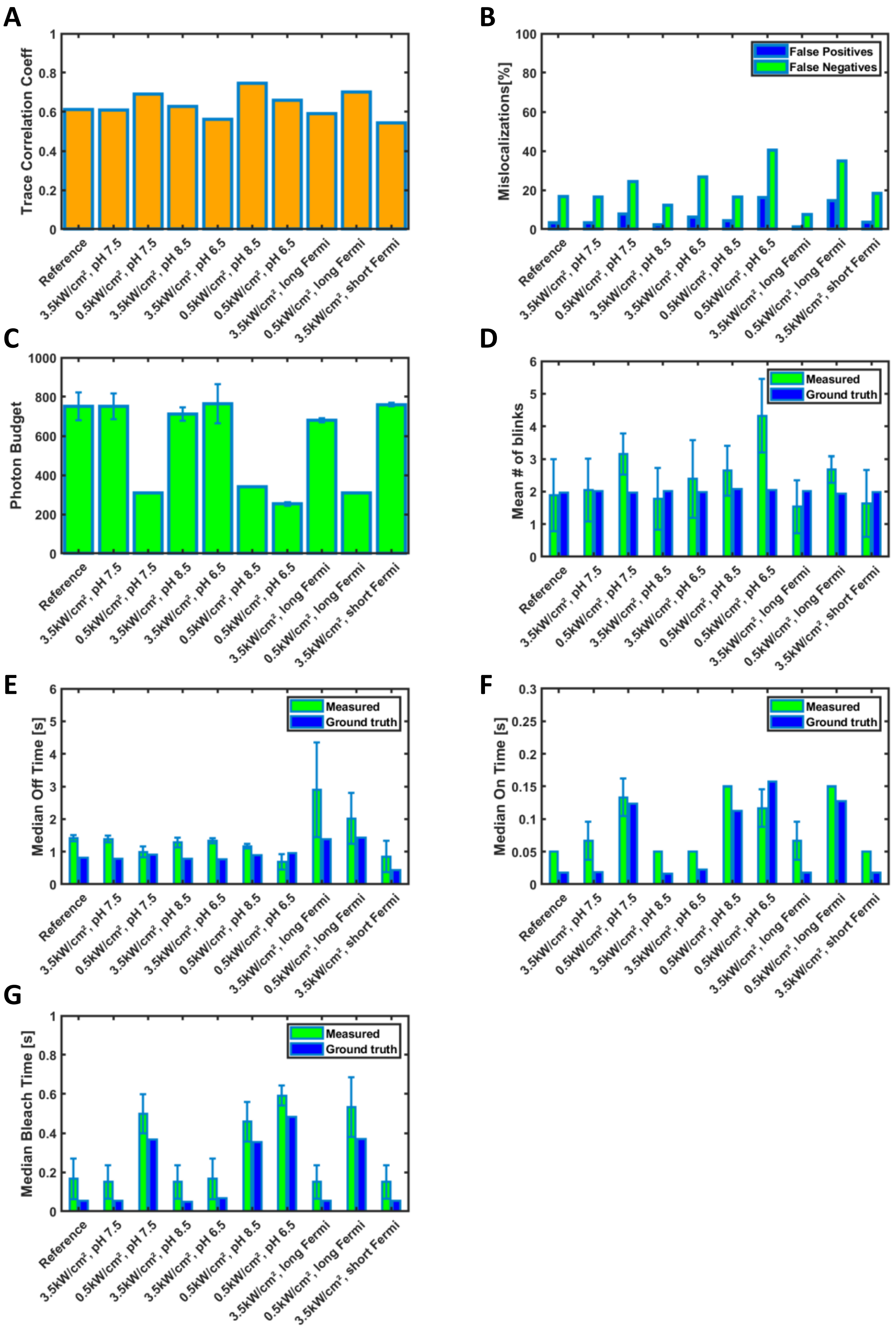


### Supplementary Fig. S4

Comparison of ground-truth versus retrieved photophysical data from SMIS simulations of Fig. 1. (A) Median fluorescence-state trace correlation; (B) False-positive (spurious localizations) and false-negative (missed localizations); (C) Median detected photon budget per localization; (D) Mean number of blinks in the photoconverted red state; (E,F,G) Median off-times, on-times and bleach time in the photoconverted red state.

Error bars in C-G where computed by splitting the dataset in three subsets along time and calculating standard deviations between the subsets.

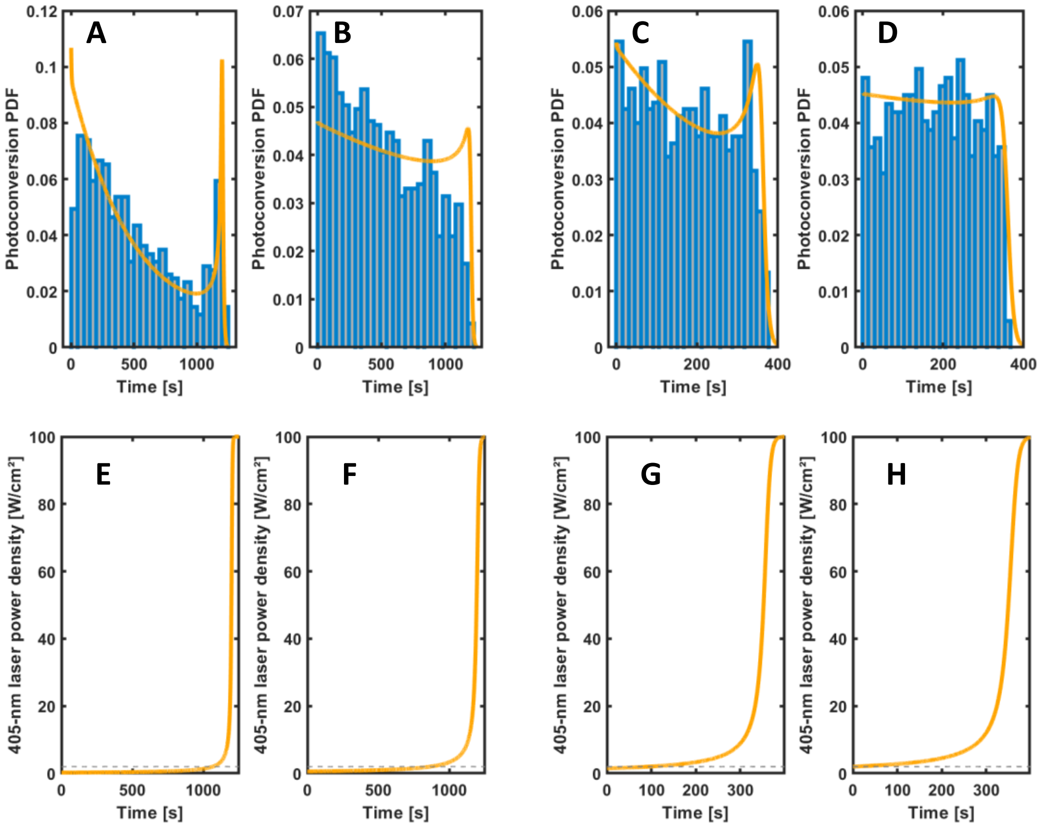


### Supplementary Fig. S5

(A-D) Overlay of theoretical Probability Density Function (PDF) of photoconversion (orange) and recovered PDF from processed SMIS data (blue) along data collection schemes using Fermi profiles. (E-H) Corresponding Fermi profiles (orange) employed for the 405-nm laser, by reference to constant laser power density (dashed gray line) used in the other NPC SMIS simulations described in this work. (A,E) 3.5 kW/cm², long Fermi (1200 s), (B,F) 0.5 kW/cm², long Fermi (1200 s), (C,G) 3.5 kW/cm², short Fermi (400 s), (D,H) 0.5 kW/cm², short Fermi (400 s)

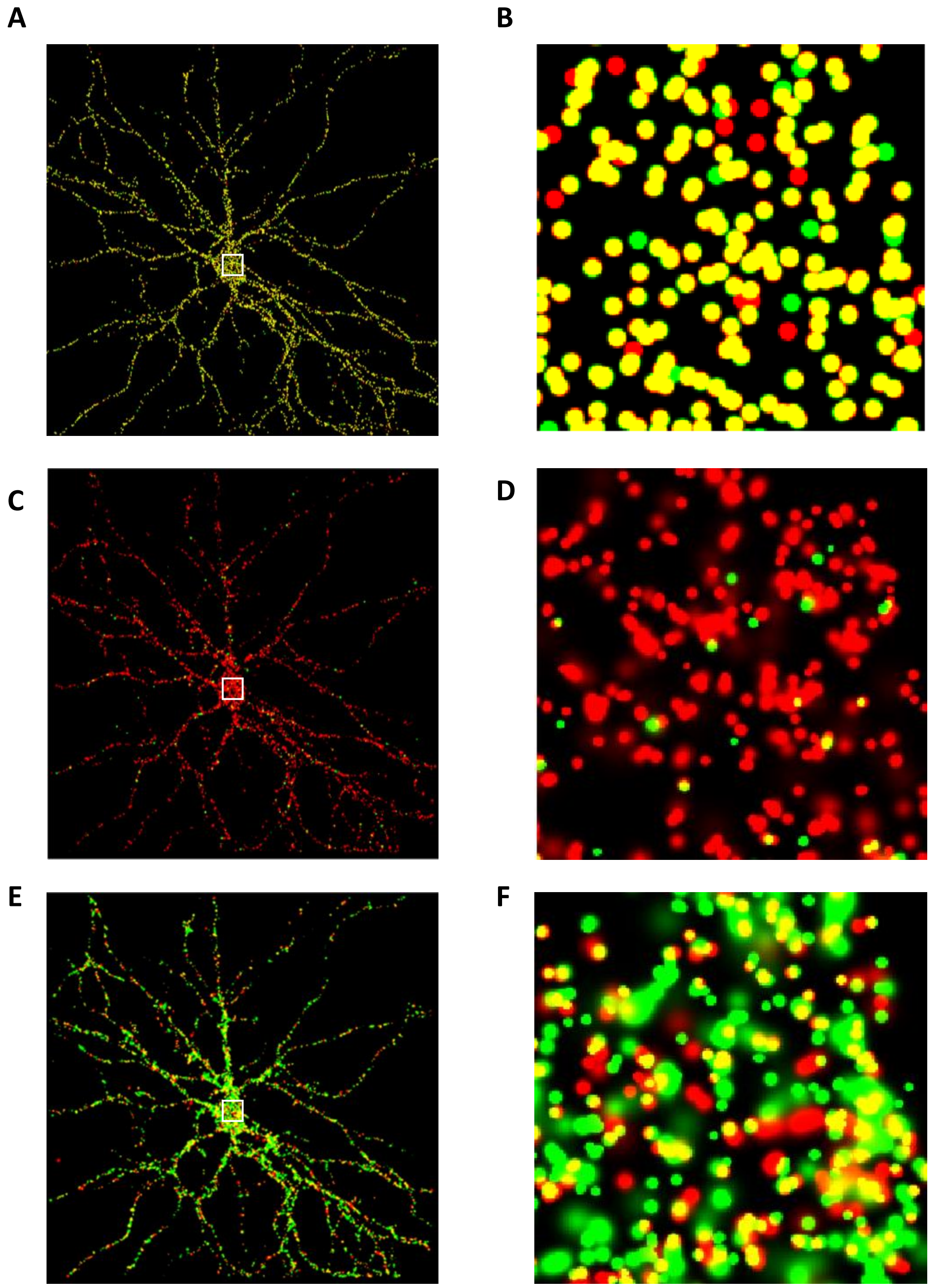


**Supplementary Fig. S6**

Pseudo-2 color SMIS simulations related to Fig. 2. (A-B) Ground truth colocalization of mEos4b (green) and Dendra2 (red) (C-D) Colocalization of mEos4b (green, in absence of mEos4b green-state photoswitching) and Dendra2 (red), (E-F) Colocalization of PAmCherry (green) and Dendra2 (red). Areas within the white squares in (A,C,E) are enlarged in (B,D,F) for inspection of colocalization events.

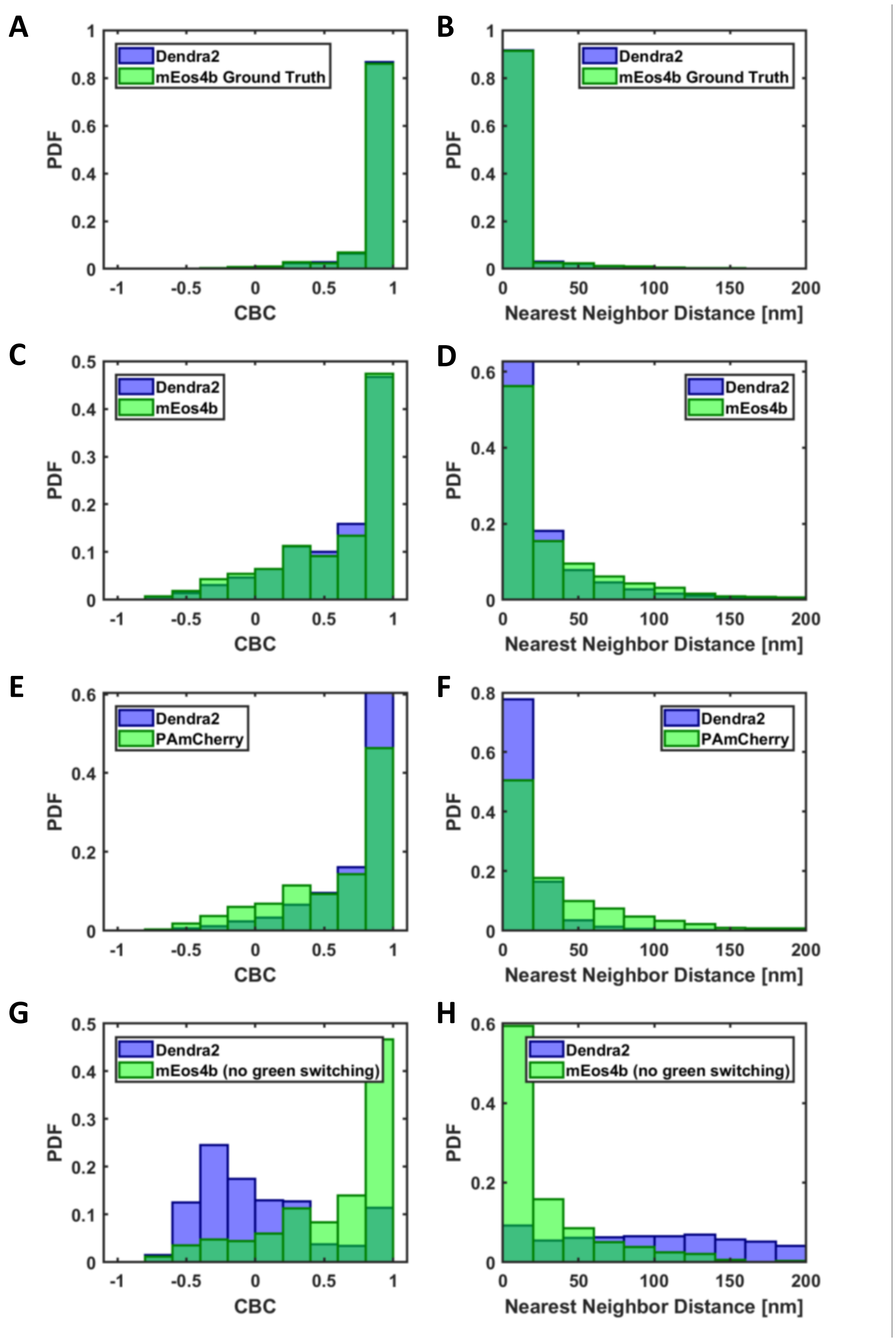


### Supplementary Fig. S7

Histograms of coordinate based colocalizations (CBC) indices and nearest neighbor distances for the various fluorescent protein pairs tested. (A-B) Ground truth colocalization of mEos4b and Dendra2. (C-D) Measured colocalization of mEos4b and Dendra2. (E-F) Measured colocalization of PAmCherry and Dendra2. (G-H) Measured colocalization of mEos4b (no green state photoswitching) and Dendra2. PDF : probability density function.

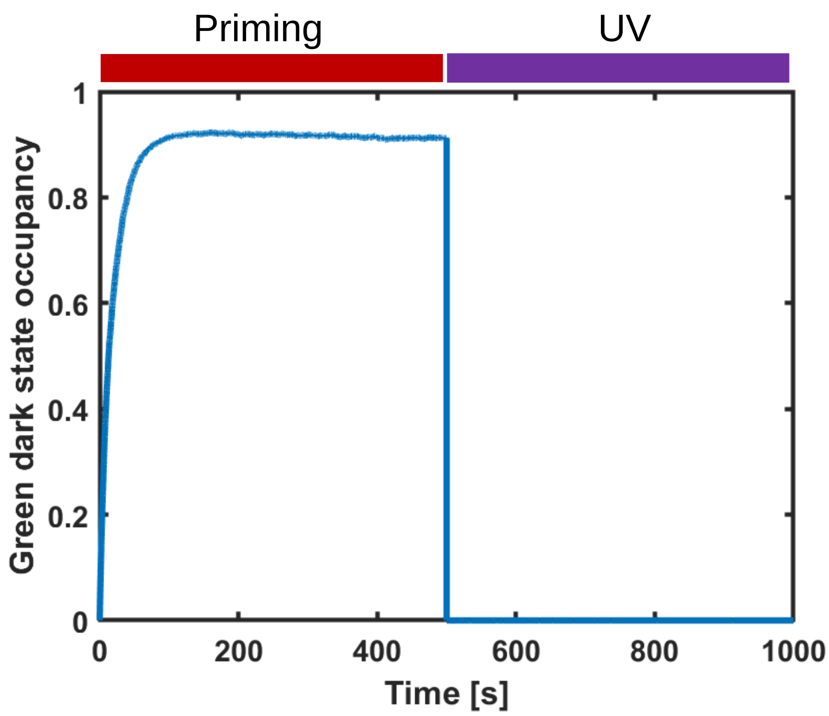


**Supplementary Fig. S8**

Occupancy of the mEos4b long-lived dark state during pseudo-2 color SMIS data collection (priming phase followed by UV phase).

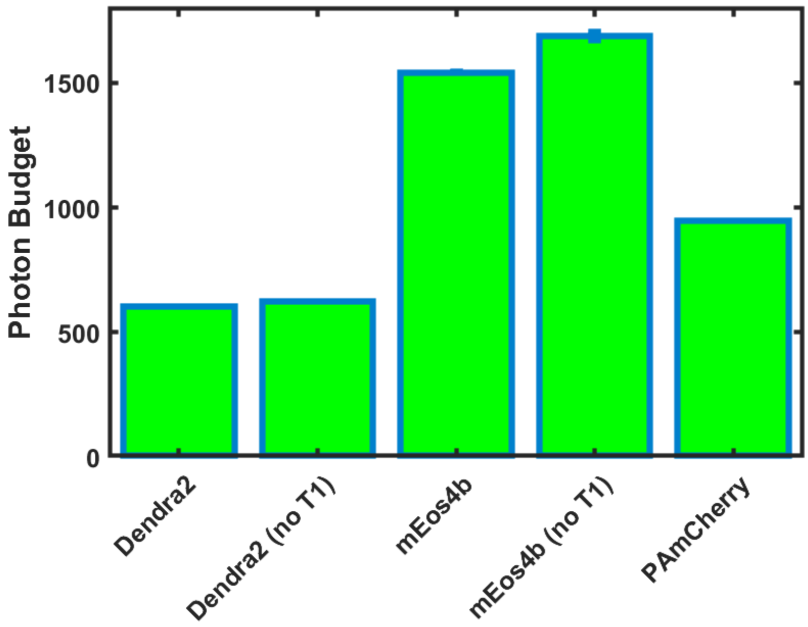


**Supplementary Fig. S9**

Photon-budget per localization during pseudo-2 color SMIS data collection. In Dendra2 (no T1) and mEos4b (no T1), intersystem crossing of the red chromophore to the triplet state was artificially suppressed, resulting in higher photon budgets per localization.

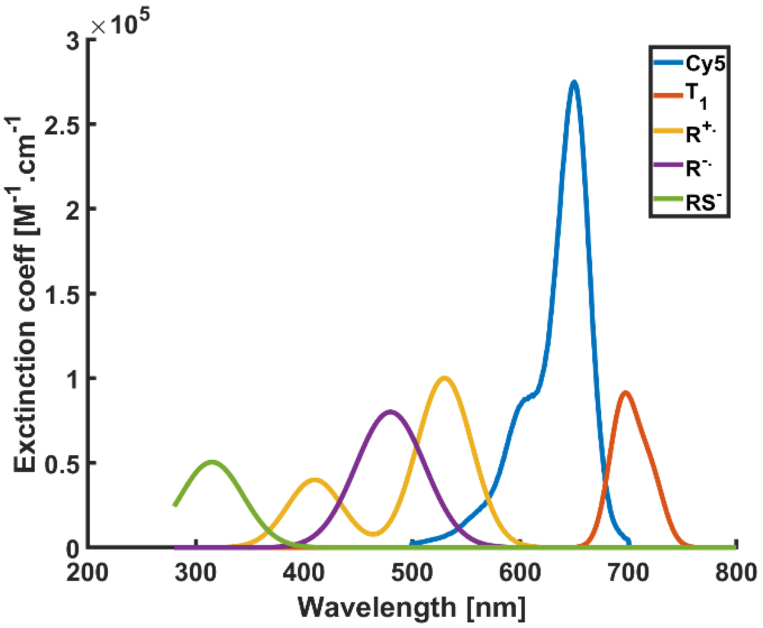


### Supplementary Fig. S10

Absorption spectra of Cy5 photophysical states used in dSTORM SMIS simulations of Fig. 3. Spectra were generated with the dedicated SMIS tool, based on spectral data of Fig. 3 in ref 20.

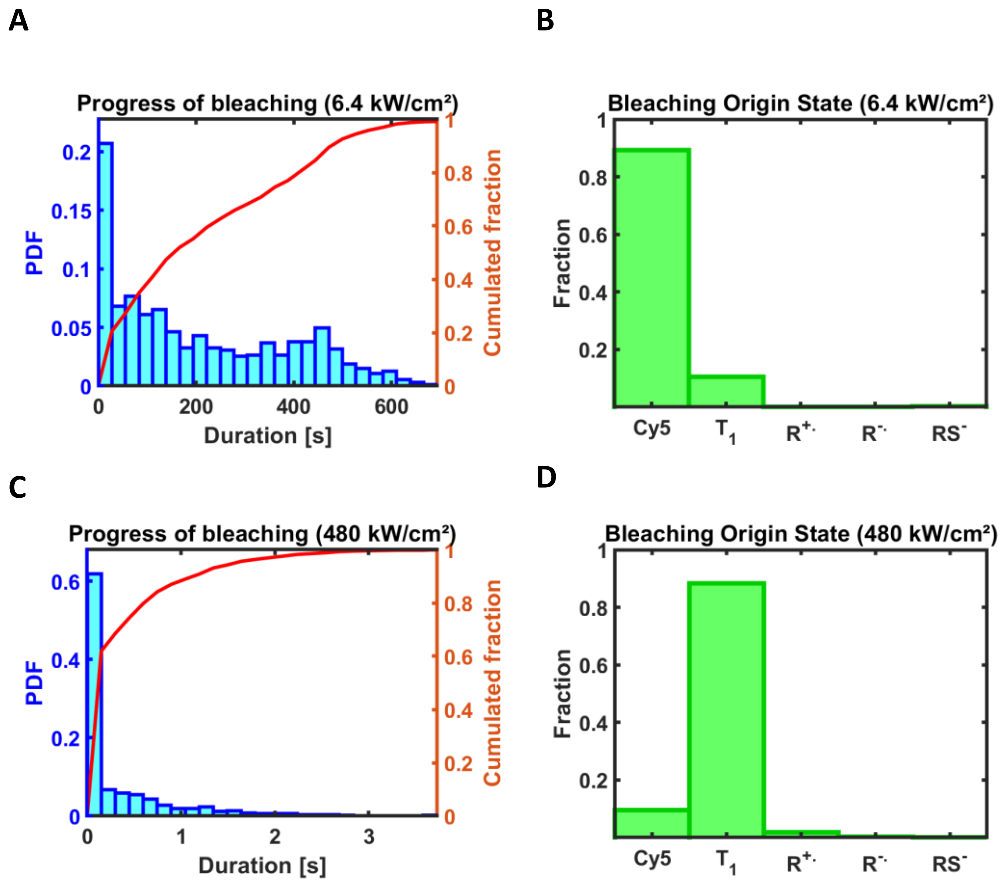


### Supplementary Fig. S11

Progress of Cy5 photobleaching (A,C) and photophysical state at the origin of bleaching (B,D) with 6.4 kW/cm $^2$ (A,B) or 480 kW/cm$^2$ (C,D) 647 nm laser power density. PDF : probability density function.

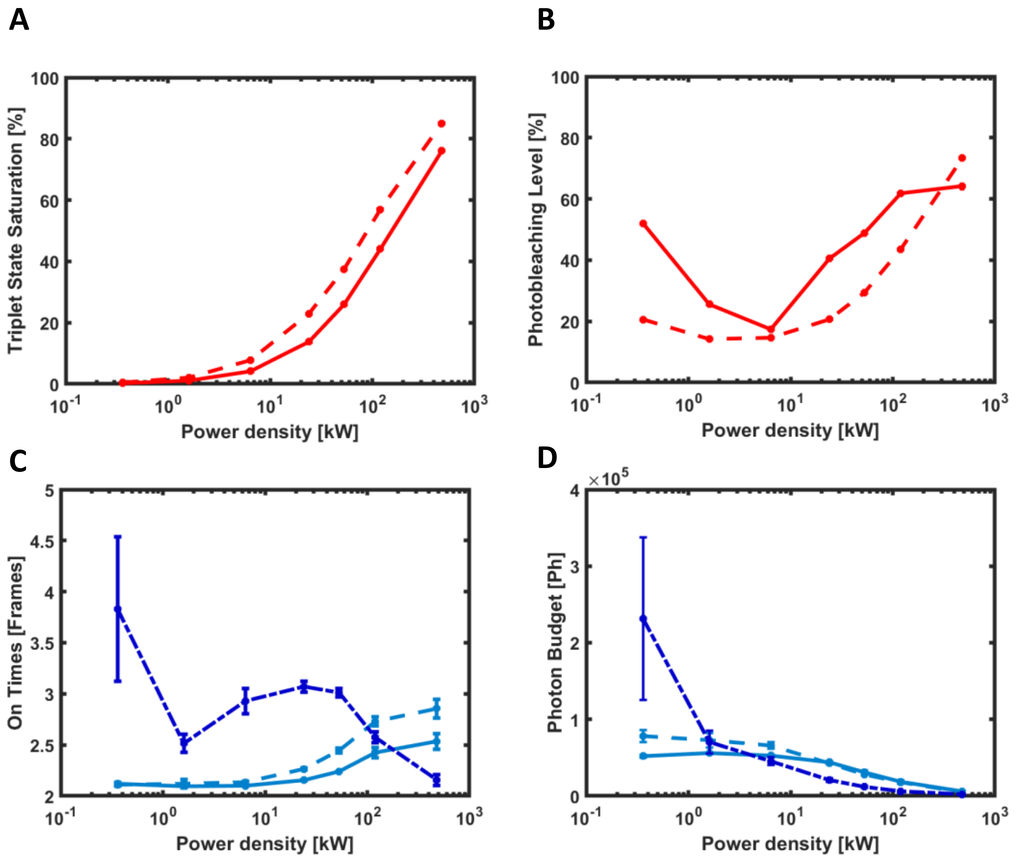


### Supplementary Fig. S12

Triplet-state saturation level of Cy5 (A) and photobleaching level reached after illumination for 1000 frames (B), as a function of 647 nm light power density, using photophysical parameters of ref 20 (plain lines) or corrected parameters (dashed line). On times (C) and photon budget (D) from experimental data from ref 11 (dark blue dashed line), or from SMIS simulations using photophysical parameters of ref 20 (light blue plain lines) or corrected parameters (light blue dashed line). Error bars for SMIS data show standard deviation for n=3 simulations.

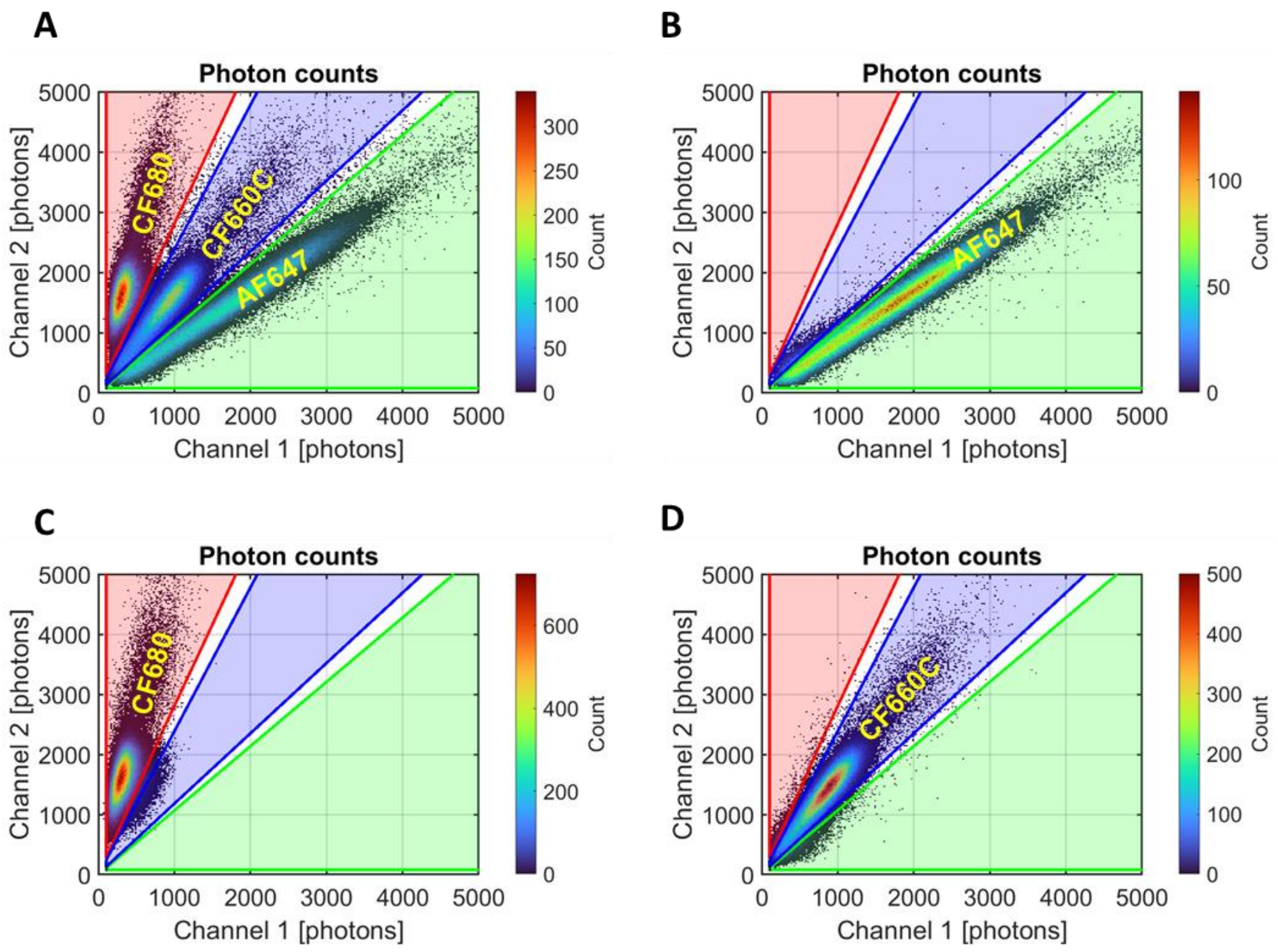


**Supplementary Fig. S13**

Bivariate histograms of photon counts for spectral demixing of AF647, CF660C and CF680. The numbers of fluorophores with specified numbers of detected photons in the two channels (channel 1: shorter wavelengths, channel 2: longer wavelengths) are shown according to color maps displayed on the right of the plots. Regions of interest (ROIs) for unmixing are shown in red, green and blue. (A) 3-color dataset. (B) control for AF647. (C) control for CF680. (D) control for CF660C

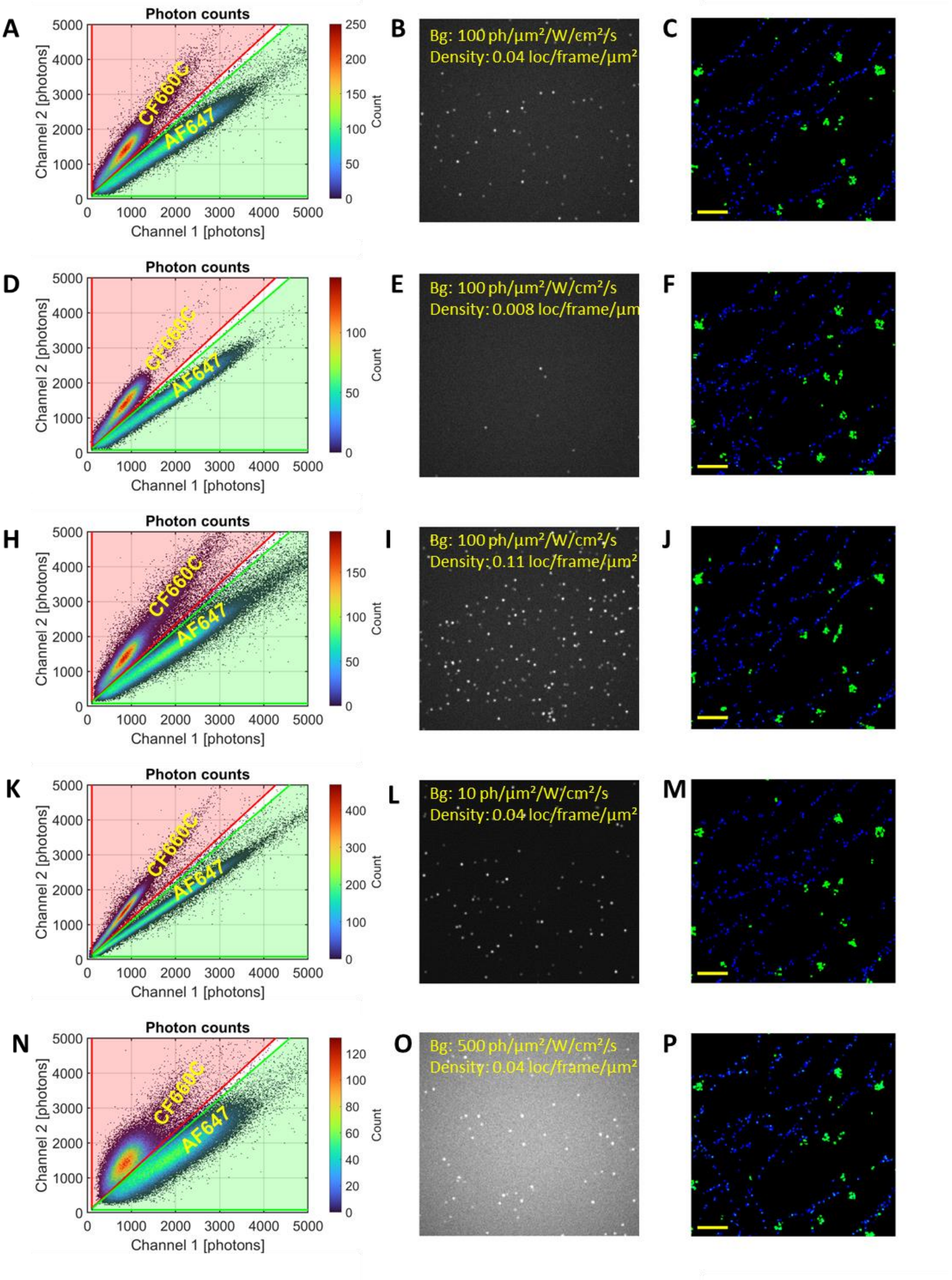


### Supplementary Fig. S14

Sensitivity of spectral demixing to localization density and fluorescence background in 2-color experiments with AF647 and CF660C. From top to bottom: medium background/ medium localization density; medium background/ low localization density; medium background/ high localization density; low background/ medium localization density; high background/ medium localization density. (A,D,H,K,N): Bivariate histograms of photon counts. The numbers of fluorophores with specified numbers of detected photons in the two channels (channel 1: shorter wavelengths, channel 2: longer wavelengths) are shown according to color maps displayed on the right of the plots. Regions of interest (ROIs) for unmixing are shown in red and green. (B,E,I,L,O) raw SMIS frame (frame #100) acquired in channel 1 for each condition indicated in yellow. Bg: background fluorescence, Density: localization density. (C,F,J,M,P) extracts of rendered 2-color images for each condition. Blue: AF647; Green: CF660C. Scale bar: 1 μm.

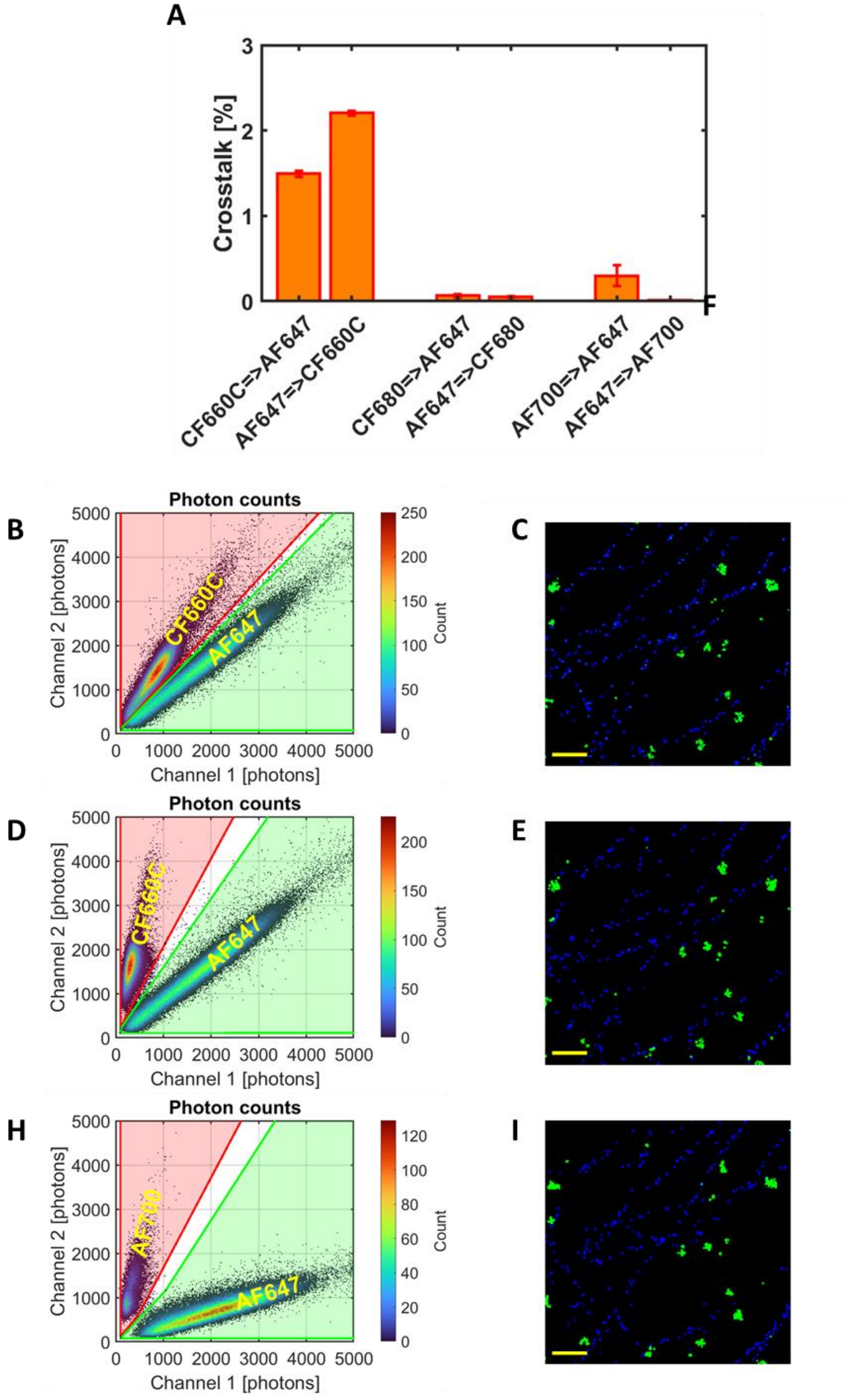


### Supplementary Fig. S15

Sensitivity of spectral demixing to fluorophore photophysics in 2-color experiments with AF647 in tandem with CF660C, CF680 or AF700. (A) Crosstalk between pairs of fluorophores. Error bars were computed by randomly splitting simulated datasets in 3 subsets. (B,D,H): Bivariate histograms of photon counts. The numbers of fluorophores with specified numbers of detected photons in the two channels (channel 1: shorter wavelengths, channel 2: longer wavelengths) are shown according to color maps displayed on the right of the plots. Regions of interest (ROIs) for unmixing are shown in red and green. (C,E,I) extracts of rendered 2-color images. Blue: AF647; Green: CF660C (C), CF680 (E), AF700 (I). Scale bar: 1 μm.

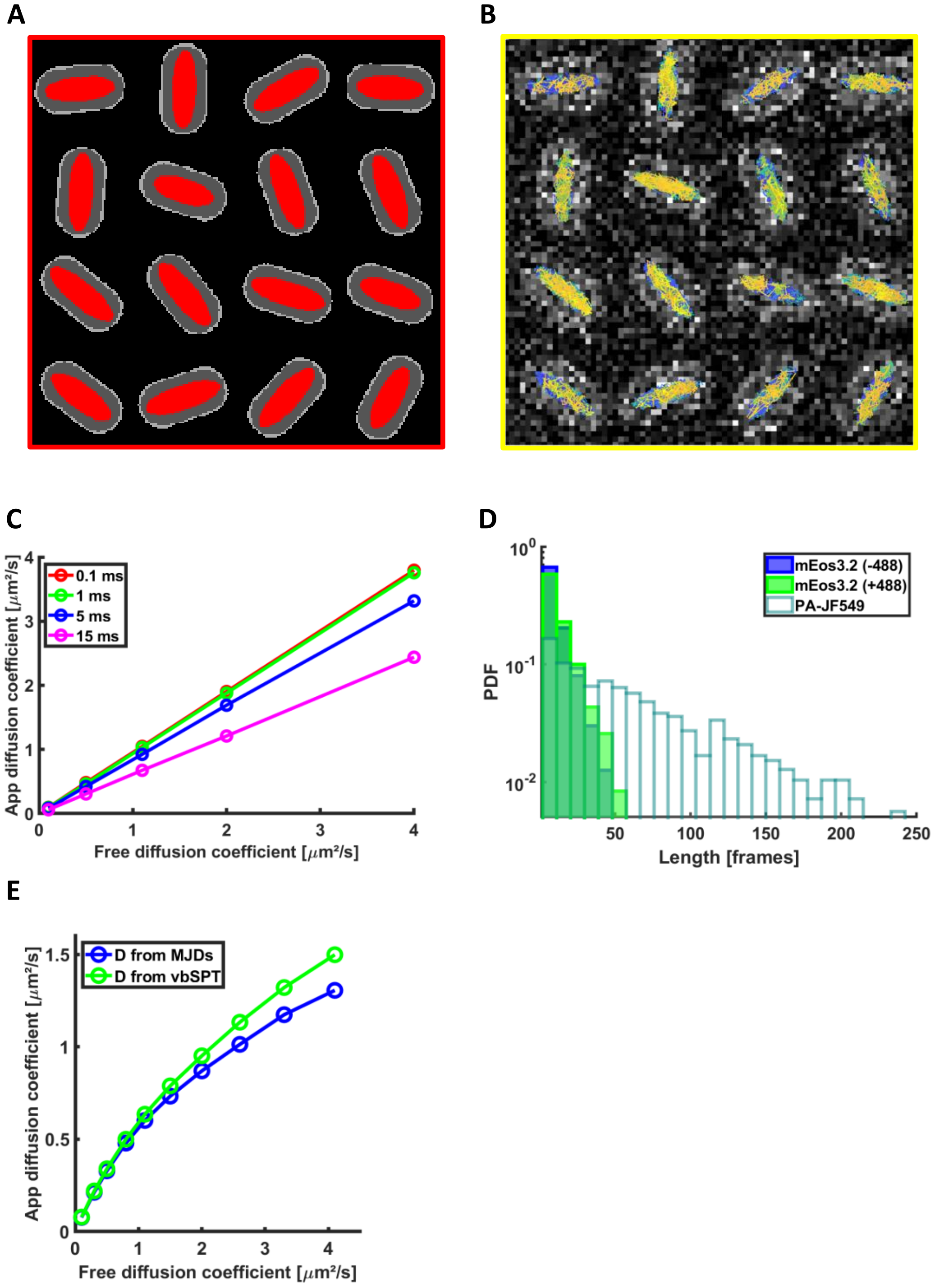


### Supplementary Fig. S16

(A) Extract of the bacterial field used for the sptPALM simulations of Fig. 4, highlighting in red the nucleoid in which the labeled NAP diffuse. (B) Same area of the bacterial field virtual sample highlighting mEos3.2 tracks superimposed on a SMIS-generated CCD frame showing autofluorescence produced by individual bacteria. Figure produced with Trackit (ref 26). (C) Effect of temporal averaging on non-confined diffusing mEos3.2

molecules. Input free diffusion coefficients are shown on the X-axis, whereas apparent diffusion coefficients extracted from MJD histograms are shown on the Y-axis. Apparent diffusion coefficients are smaller when progressively increasing CCD exposure while progressively decreasing dark times between frames, at constant total frame time (15 ms). (D) Tracklength histogram of mEos3.2 in the absence or presence of additional 488 nm light, compared to PA-JF549 measured upon processing of the generated SMIS image stacks. (E) Calibration curves assuming single populations of fluorophores moving with increasing diffusion coefficients within the confined *E. coli* nucleoid, and showing measured apparent diffusion coefficients versus input free diffusion coefficients, at fixed frame time (5 ms CCD exposure + 10 ms dark time). PDF: probability density function

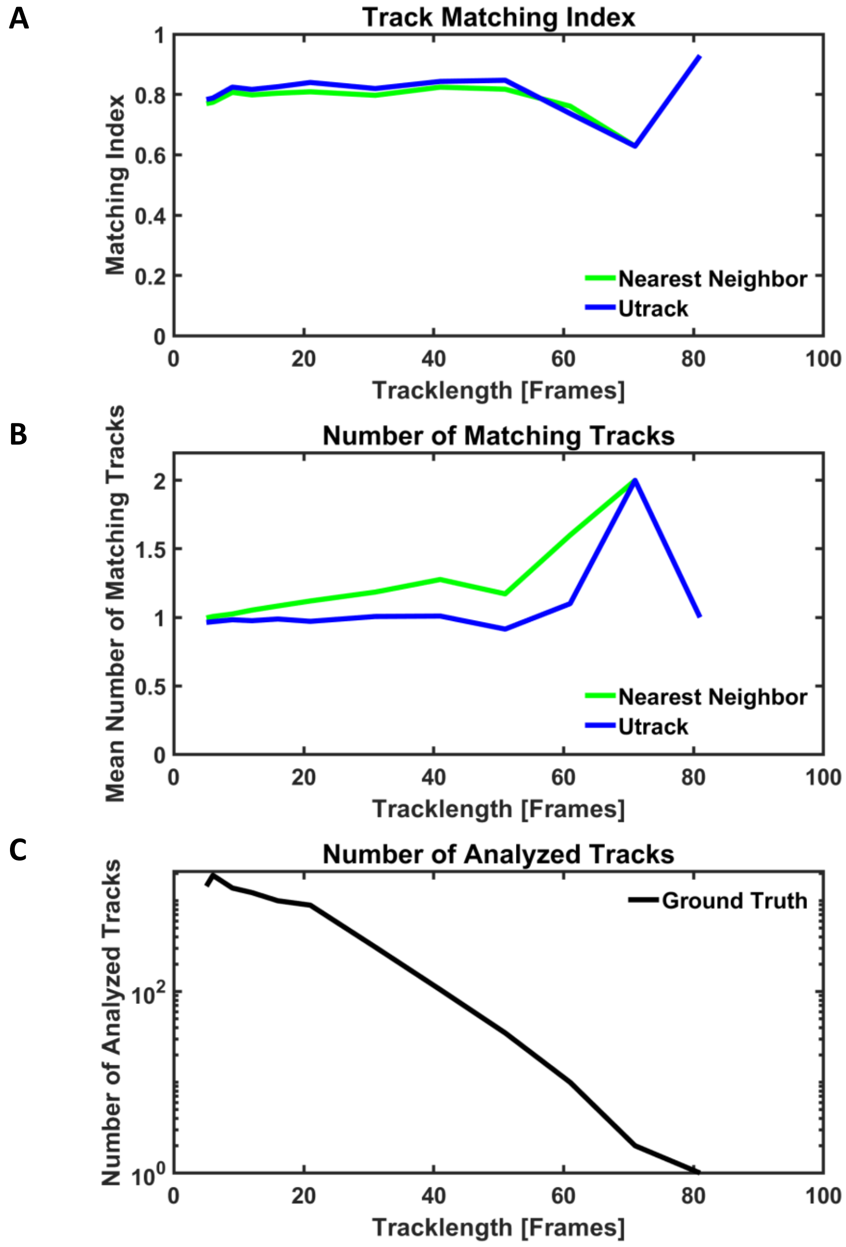


**Supplementary Fig. S17**

Example of the quality of track retrieval by Trackit (ref 26), using a nucleoid-confined mEos3.2 SPT SMIS dataset with 15 ms total frame time, and comparison between Nearest Neighbor (green) and UTrack (blue). (A) Track matching index: For each ground truth track (allowing a blinking gap of 2 frames), the closest experimental matching track is considered and the fraction of the ground truth localizations for which a processed localization is found at a distance < 30 nm is computed. (B) Number of matching tracks: for each ground truth track, the number of matching experimental tracks (i.e. for which a localization is found at a distance < 30 nm from a ground truth localization) is computed. (C) Number of ground truth analyzed tracks.

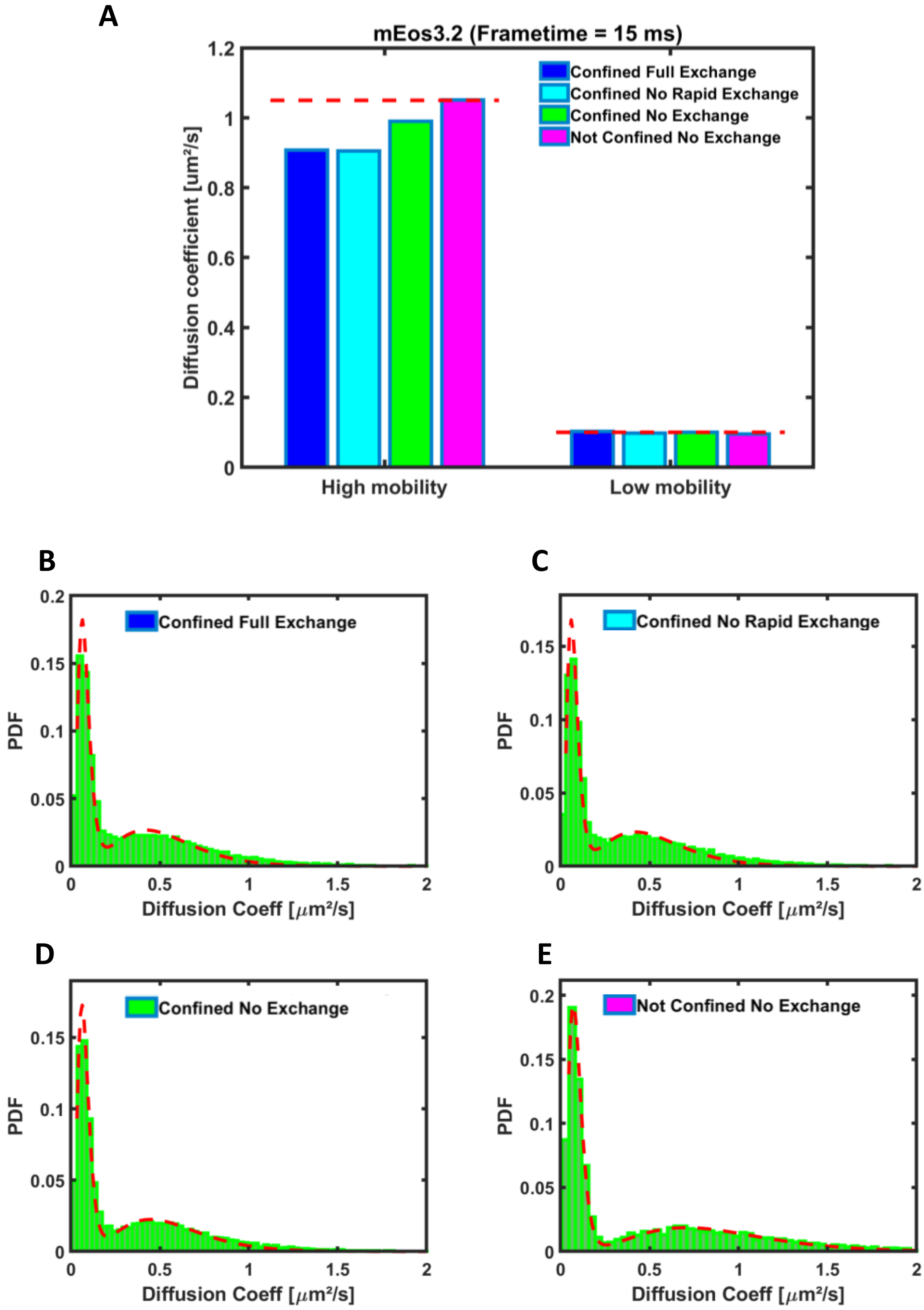


### Supplementary Fig. S18

Effect of kinetic exchange and confinement on retrieved unbiased diffusion coefficients from mEos3.2 SPT data (in the absence of 488 nm light) extracted from MJD-based diffusion coefficient histograms (tracklength = 5, total frametime = 15 ms). (A) Retrieved unbiased diffusion coefficients for the low and high mobility populations. Ground truth values are marked as dashed red lines. Nucleoid confined mEos3.2 with both slow and rapid exchange as in Fig. 4A (blue); Nucleoid confined mEos3.2 with only slow exchange (cyan); Nucleoid confined mEos3.2 without exchange (green); Non-confined mEos3.2 without exchange (magenta). Associated apparent diffusion coefficient histograms (green) are shown in (B-E), together with associated fits from a 2-state model (red). The quality of the fits progressively improves from panel B to panel E.

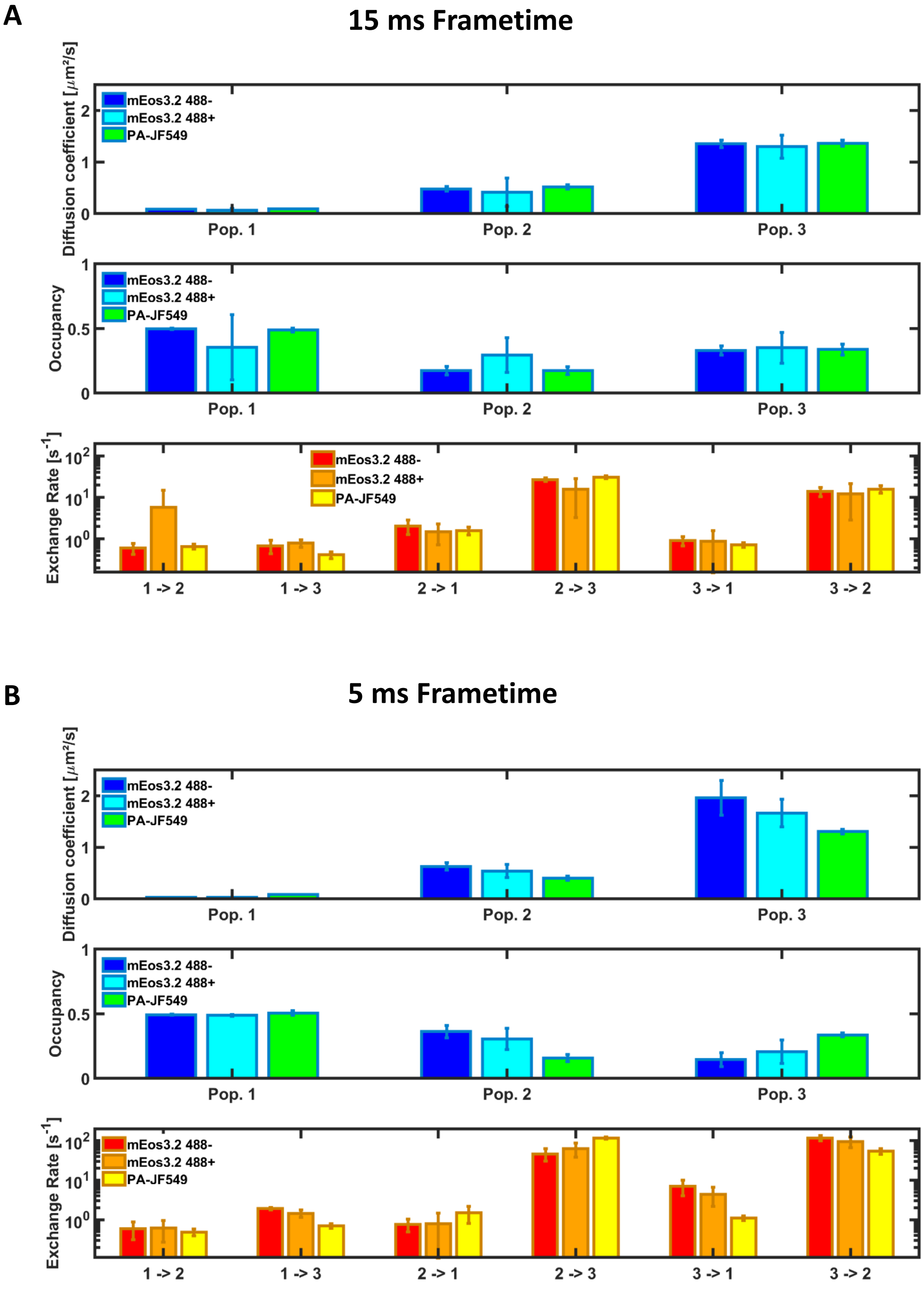


### Supplementary Fig. S19

Retrieved unbiased diffusion coefficients, population occupancies and exchange rates output by vbSPT as the most likely diffusion model. Error bars show standard deviation for n=3 simulations. (A) 15 ms total frametime. (B) 5 ms total frametime. In both cases, output values significantly deviate from ground truth values, show substantial standard deviations, and differ substantially between (A) and (B), suggesting that the 3-state models are false.

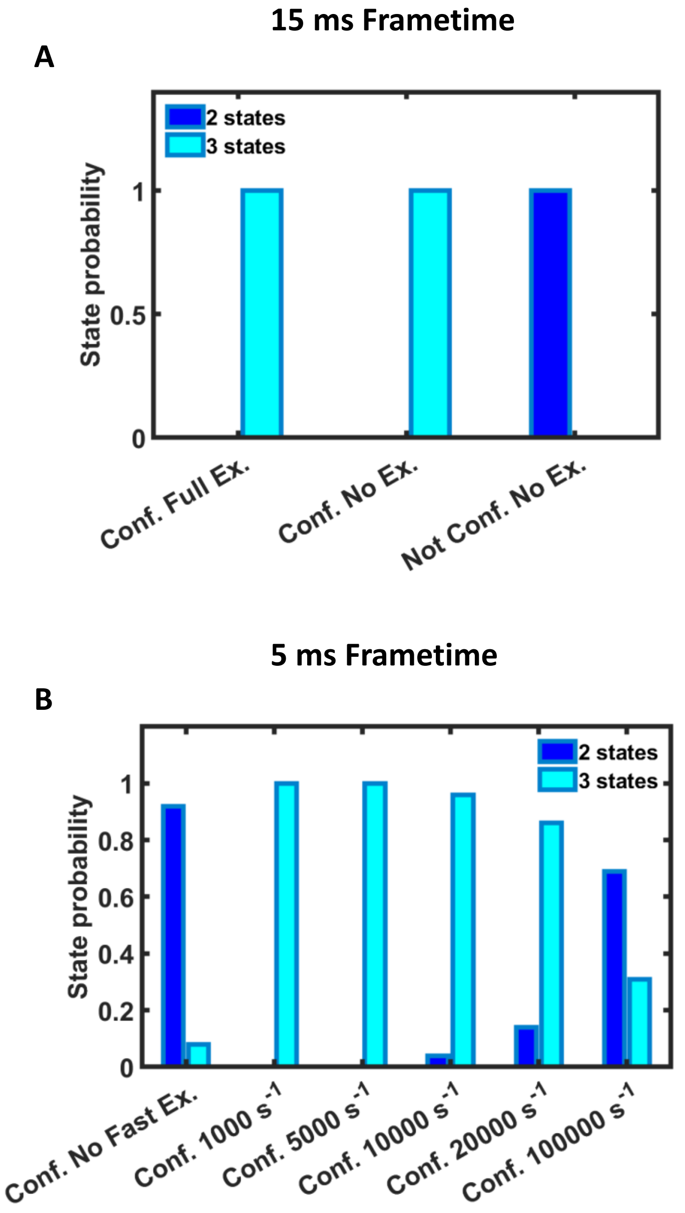


**Supplementary Fig. S20**

Probability of 2-state (dark blue) versus 3-state (light blue) models assessed by bootstrapping in vbSPT. (A) mEos3.2 SPT SMIS datasets in the absence of 488-nm light, 15 ms total frametime. Nucleoid confined mEos3.2 with both slow and rapid exchange as in Fig. 4A (Conf. Full Ex.); Nucleoid confined mEos3.2 without exchange (Conf. No Ex.); Non-confined mEos3.2 without exchange (Not Conf. No Ex.). At 15 ms total frametime, nucleoid confinement is sufficient to produce a false 3-state model. (B) PA-JF549 SPT SMIS datasets, 5 ms total frametime. The case of nucleoid confined PA-JF549 only slow exchange (Conf. No Fast Ex.) is compared to cases with both slow and rapid exchange from 1000 $s^{-1}$ to 100000 $s^{-1}$. A 2-state model is favored only in the absence of rapid exchange, or in the presence of very fast exchange.

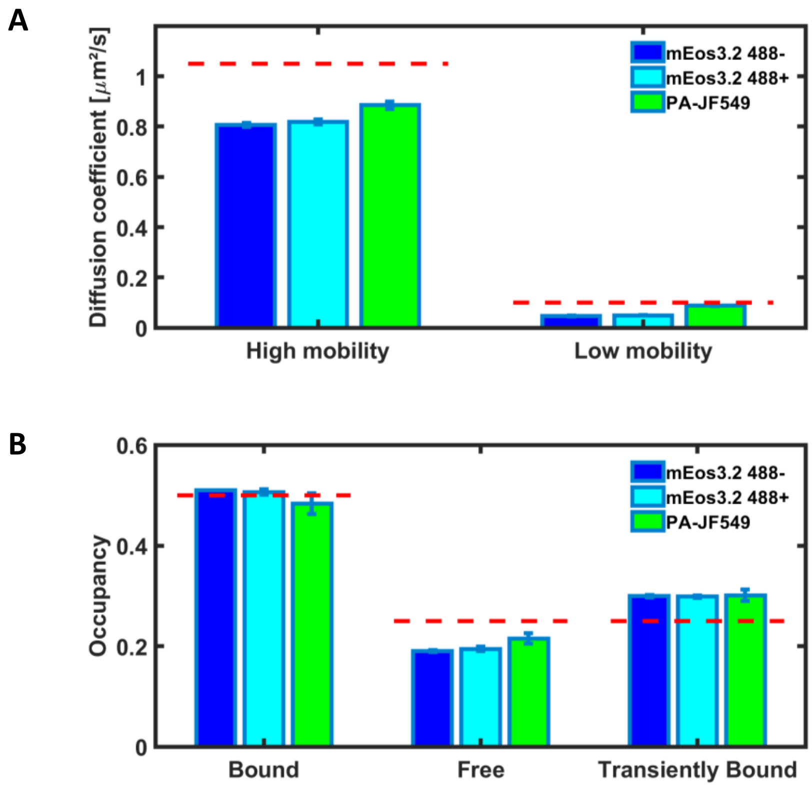


**Supplementary Fig. S21**

Unbiased diffusion coefficients (A) and diffusion state's occupancies (B) recovered from MJD's-derived diffusion coefficients for mEos3.2 in the absence (blue) or presence (cyan) of 488 nm light, and for PA-JF549 (green). Ground truth values are shown as dashed red lines. Total frame time = 5 ms, to be compared with Fig. 4D.

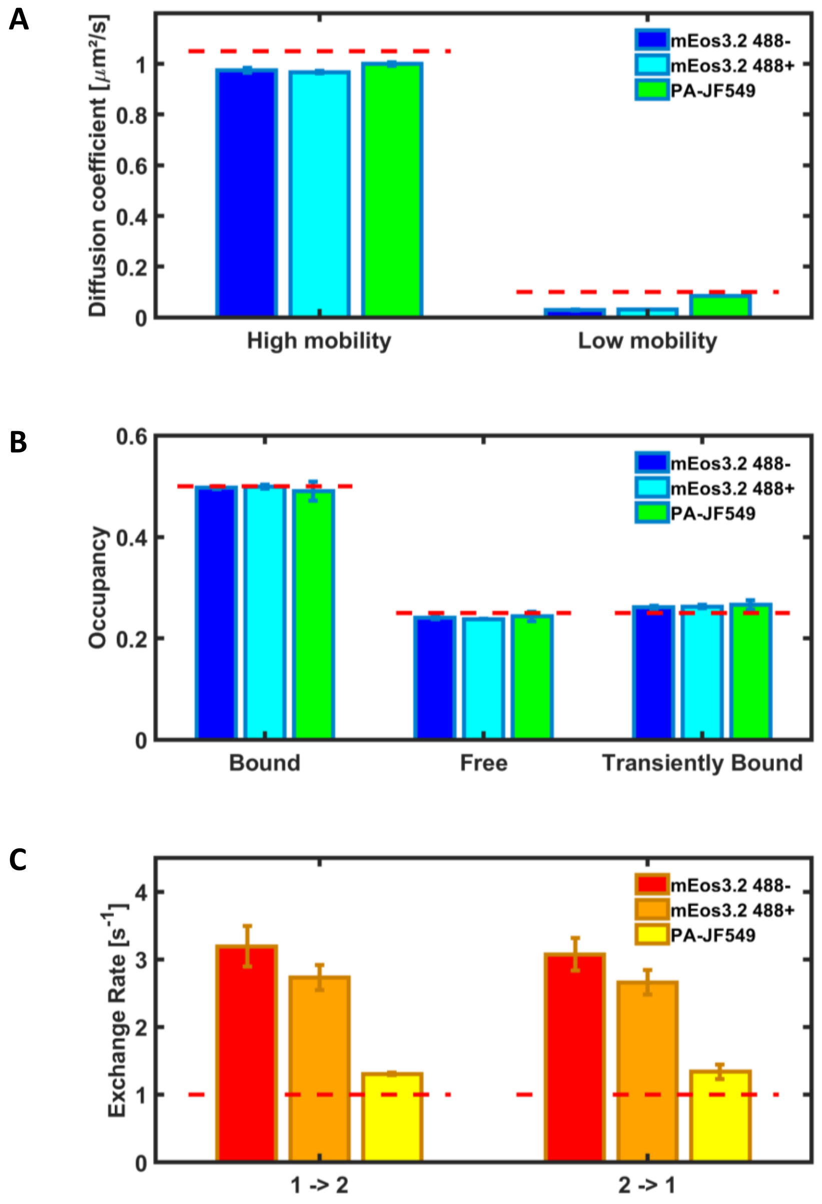


### Supplementary Fig. S22

Unbiased diffusion coefficients (A) and diffusion state's occupancies (B) recovered from vbSPT-derived diffusion coefficients for mEos3.2 in the absence (blue) or presence (cyan) of 488 nm light, and for PA-JF549 (green). (C) Slow exchange rates recovered from vbSPT, for mEos3.2 in the absence (red) or presence (orange) of 488 nm light, and for PA-JF549 (yellow). 1 → 2: Tight DNA-bound state to searching state; 2 → 1: Searching state to tight DNA-bound state. Total frame time = 5 ms, to be compared with Fig. 4E-F.

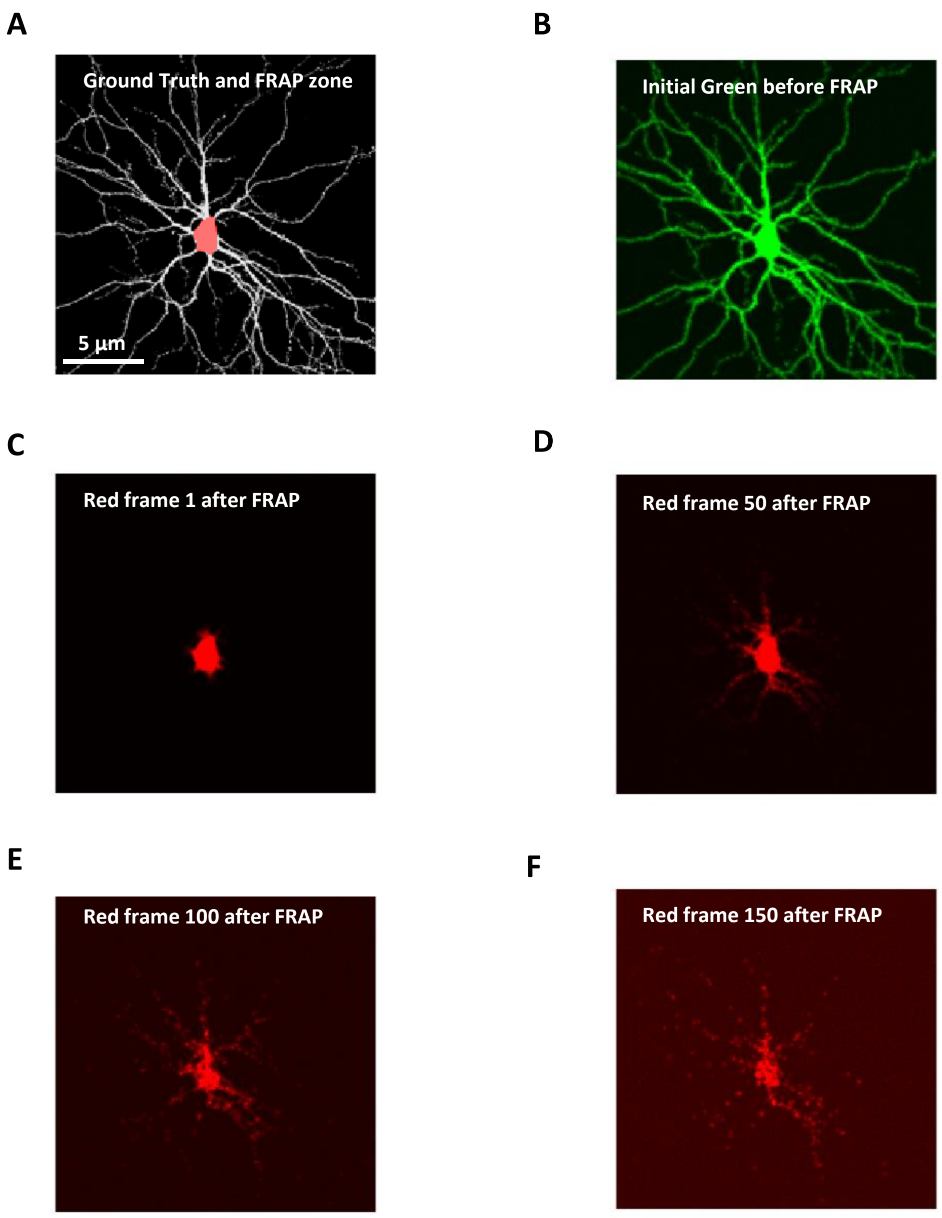


### Supplementary Fig. S23

SMIS pulse-chase experiment with mEos4b. (A) Virtual 2D neuronal cell (grey) and photoconversion zone (pink) applied at the level of the cell nucleus. (B) Initial mEos4b green image before FRAP (green channel of the virtual microscope). (C-F) mEos4b red images (red channel of the virtual microscope) immediately after photoconversion in the FRAP zone (C) and 2.5 s (D), 5 s (E) and 7.5 s (F) after photoconversion. Diffusion of the photoconverted mEos4b red molecules is clearly visible, together with progressive photobleaching. Frametime = 50 ms ; mEos4b diffusion coefficient : 2 $\mu m^2/s$. 561-nm laser power density : 200 $W/cm^2$ ; 488-nm laser power density : 100 $W/cm^2$ (1$^{st}$ frame) ; 405-nm laser power density : 5000 $W/cm^2$ (2$^{nd}$ frame, FRAP zone).

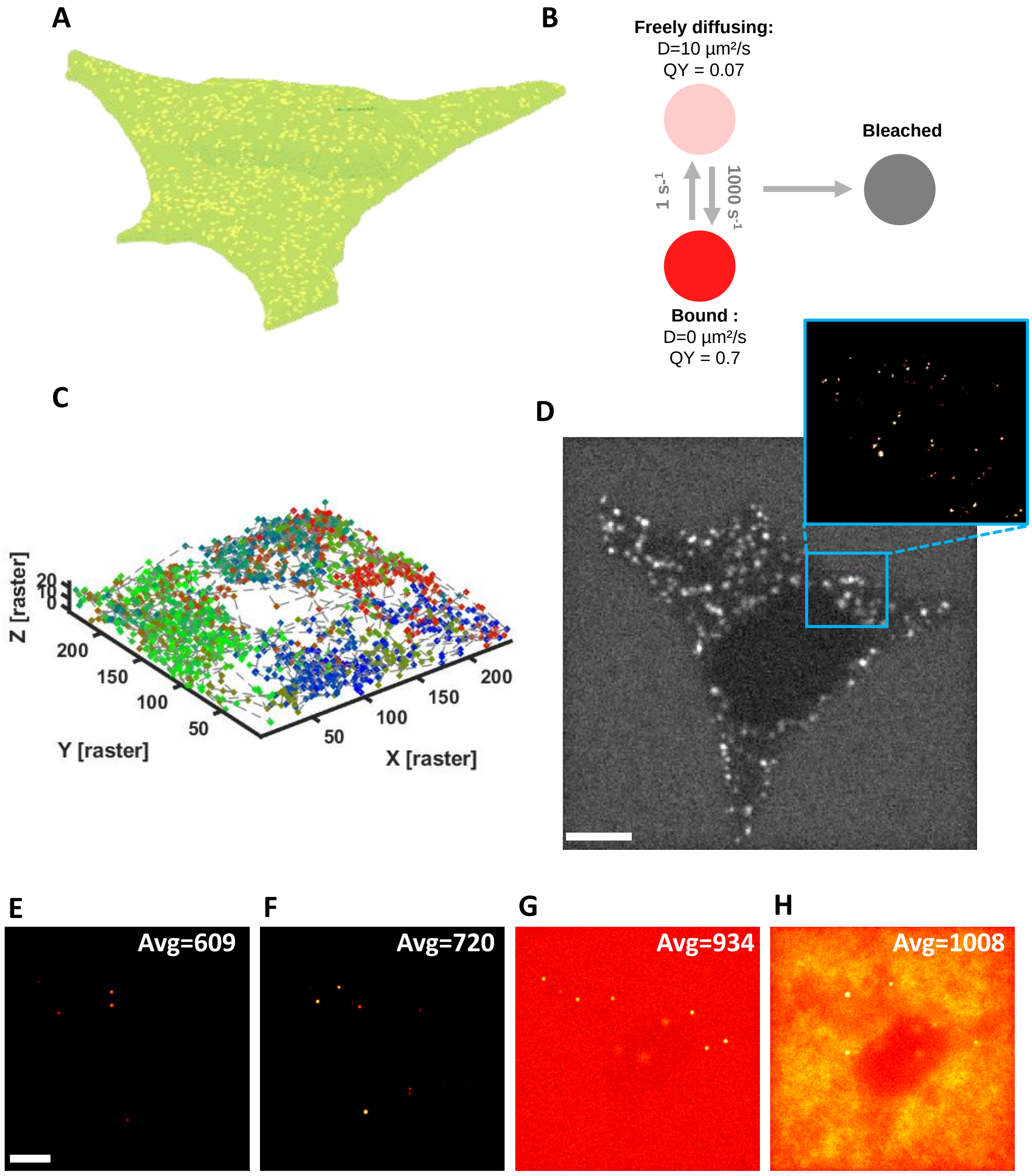


### Supplementary Fig. S24

SMIS 3D-PAINT imaging of receptor clusters anchored in the cytoplasmic membrane of a HeLa cell, using fluorogenic NileRed and HILO illumination mode. (A) 3D view of the virtual HeLa cell. Receptor clusters are drawn in yellow. (B) Employed NileRed photophysical and diffusion model (QY = fluorescence quantum yield). (C) Exemple of a single NileRed molecule diffusing around the cell for ~1800 frames (from blue to red). (D) Diffraction limited average image of the 10000 recorded frames and view of PAINT reconstructed receptor clusters (zoom from blue square in C). Scalebar = 5 µm. (E-H): Average images of 100-frames PAINT datasets. Average image counts are shown. Images are shown on the same scale. Scalebar = 5 µm. (E) Hilo mode, fluorogenic labels; (F) Hilo mode, non-fluorogenic labels. Fluorescence from the non-bound labels rapidly diffusing in the HILO zone slightly increases the average background fluorescence; (G) Widefield mode, fluorogenic labels; (H) Widefield mode, non-fluorogenic labels. Fluorescence from the non-bound labels rapidly diffusing in 3D is clearly visible and further increases the average background fluorescence. Frametime = 50 ms. 561-nm laser power density : 200 W/cm². Autofluorescence : 300 ph/pixel/s.

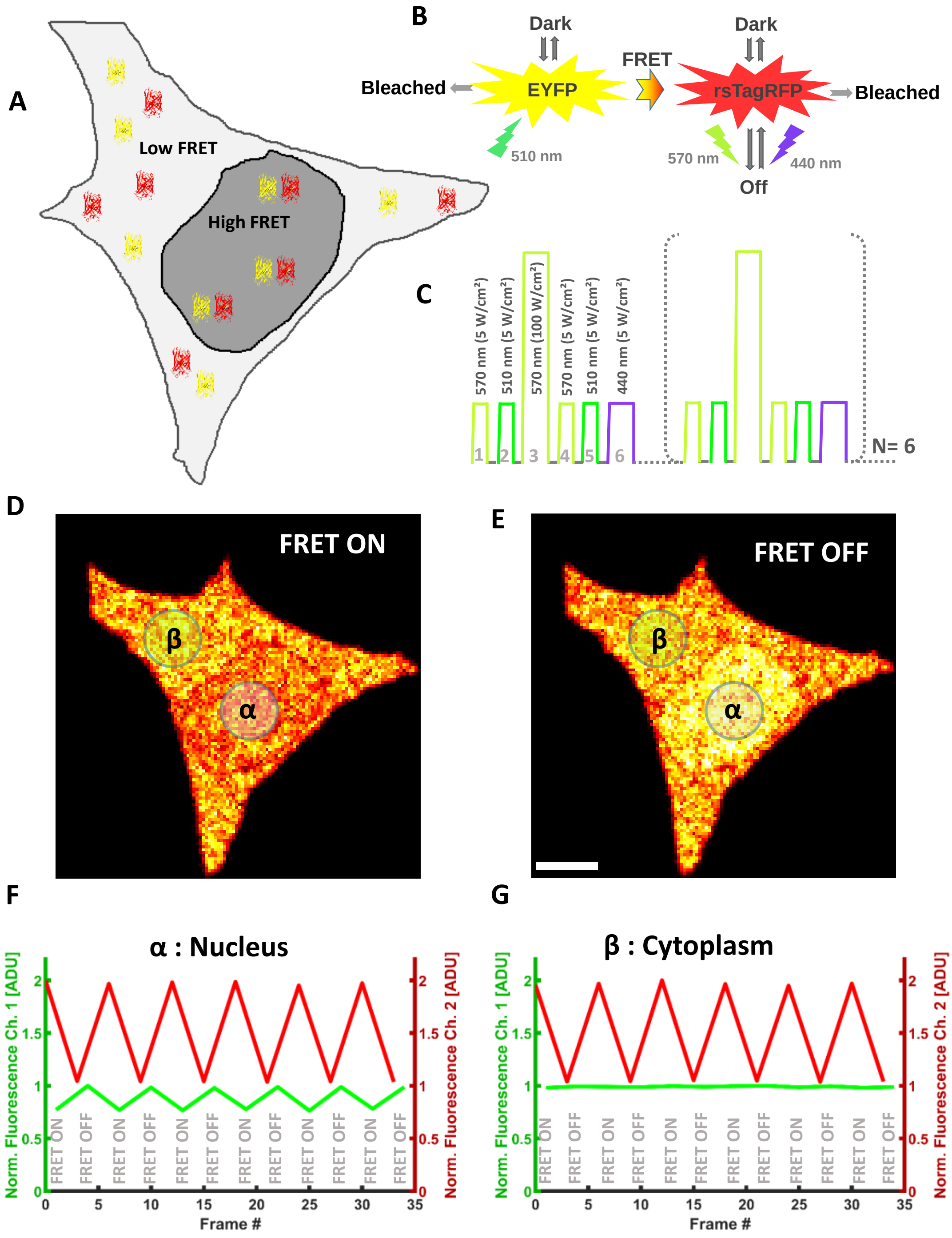


### Supplementary Fig. S25

SMIS photochromic FRET experiment with EYFP and rsTagRFP. (A) 2D view of the virtual HeLa cell. EYFP and rsTagRFP colocalize in the nucleus (high FRET, distance 6 nm +/- 1 nm), but not in the cytoplasm (low FRET). (B) Employed photophysical model. Both EYFP and rsTagRFP can bleach or transiently enter a short-lived dark state that thermally recovers at 20 $s^{-1}$. rsTagRFP photoswitches under 570 nm (on to off) or 440 nm (off to on) light illumination, respectively. (C) Used laser pulse sequence : pulses 1 and 4 (570 nm) correspond to readout frames to monitor the state of rsTagRFP. Pulses 2 and 5 (510 nm) correspond to readout frames to monitor the state of EYFP. Pulse 3 (570 nm) and 6 (440 nm) are used to switch rsTagRFP off and on, respectively. (D-E) : Fluorescence image recorded in the EYFP channel from pulse 2 ((D), rsTagRFP mostly in the on-state) and 5 ((E), rsTagRFP mostly in the off-state). Regions $\alpha$ and $\beta$ delineate regions of interest from which the average recorded fluorescence intensity is plotted in F and G. (F-G) Evolution of the EYFP normalized fluorescence (green, left axis, EYFP channel) from region $\alpha$ (F) and $\beta$ (G) measured by pulses 2 (high FRET) and 5 (low FRET). The lower signal with pulses 2 is a manifestation of a FRET efficiency of ~ 20%. The evolution of the rsTagRFP normalized fluorescence (red, right axis, rsTagRFP channel, offset of 1 unit added) shows photoswitching recorded by pulses 1 (on-switched) and 4 (off-switched). Frametime = 50 ms.

A

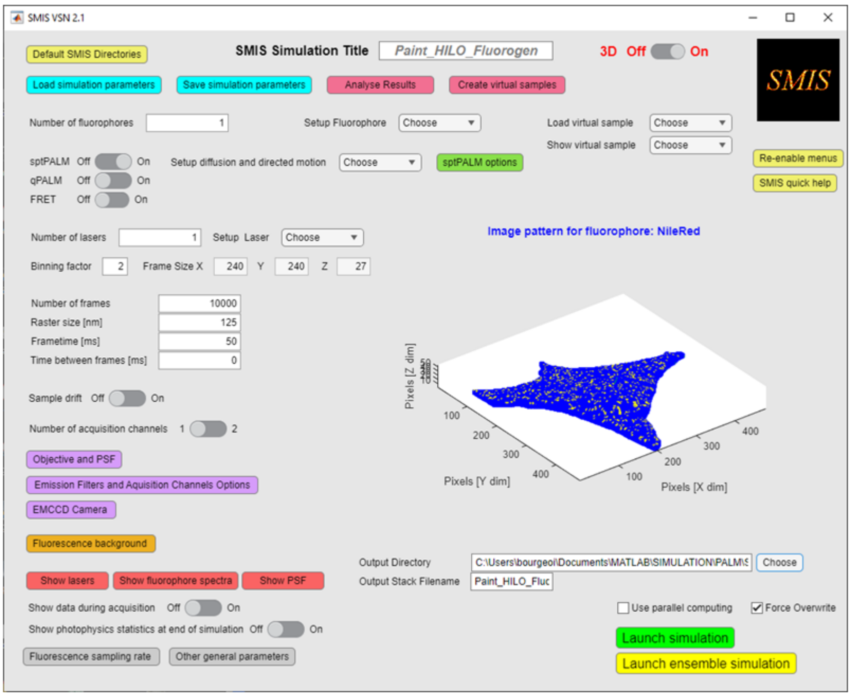


B

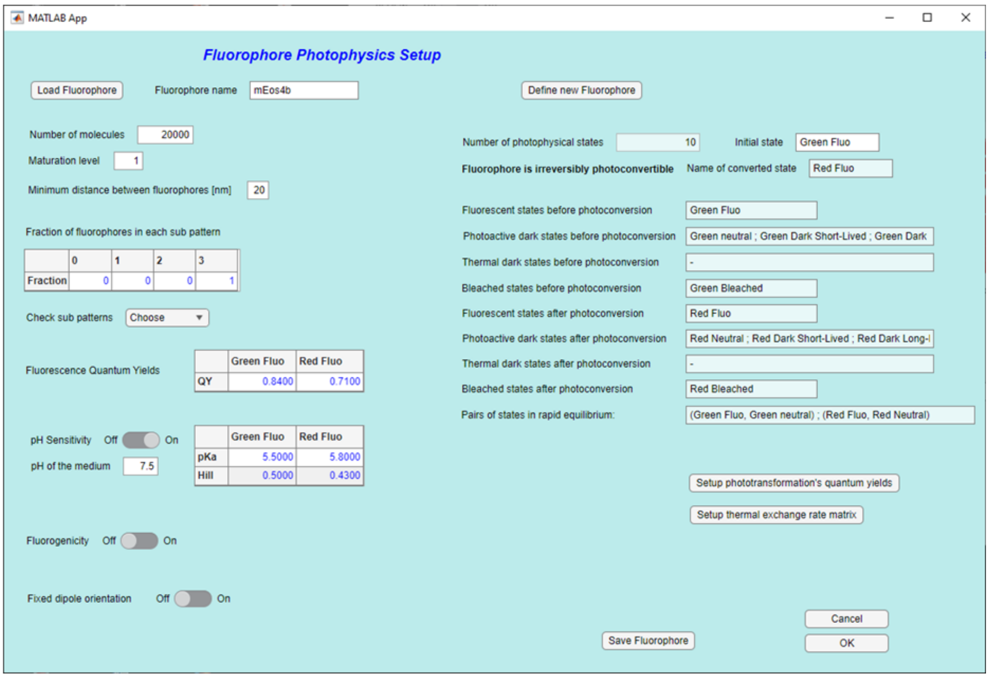


C

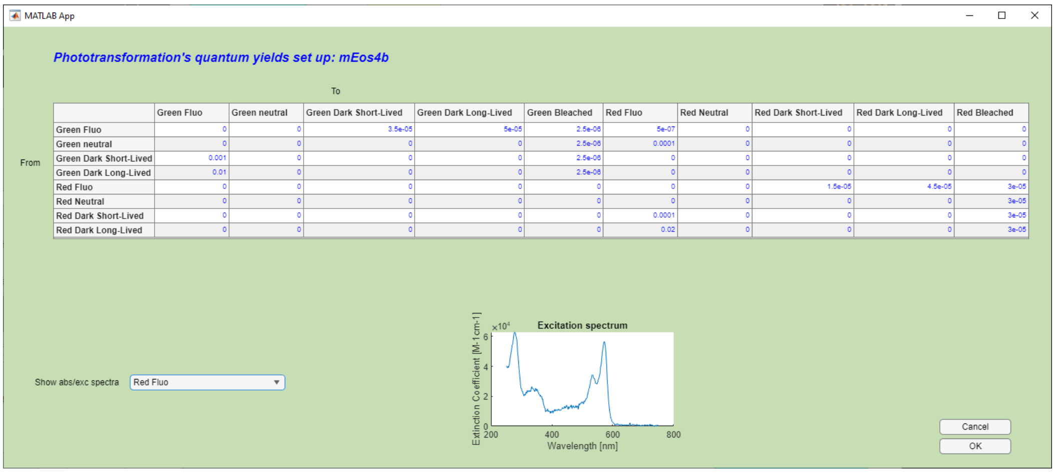


| From \ To | Green Fluo | Green neutral | Green Dark Short-Lived | Green Dark Long-Lived | Green Bleached | Red Fluo | Red Neutral | Red Dark Short-Lived | Red Dark Long-Lived | Red Bleached |
|---|---|---|---|---|---|---|---|---|---|---|
| Green Fluo | 0 | 0 | 3.5e-05 | 5e-05 | 2.5e-06 | 5e-07 | 0 | 0 | 0 | 0 |
| Green neutral | 0 | 0 | 0 | 0 | 2.5e-06 | 0.0001 | 0 | 0 | 0 | 0 |
| Green Dark Short-Lived | 0.001 | 0 | 0 | 0 | 2.5e-06 | 0 | 0 | 0 | 0 | 0 |
| Green Dark Long-Lived | 0.01 | 0 | 0 | 0 | 2.5e-06 | 0 | 0 | 0 | 0 | 0 |
| Red Fluo | 0 | 0 | 0 | 0 | 0 | 0 | 0 | 1.5e-05 | 4.5e-05 | 3e-05 |
| Red Neutral | 0 | 0 | 0 | 0 | 0 | 0 | 0 | 0 | 0 | 3e-05 |
| Red Dark Short-Lived | 0 | 0 | 0 | 0 | 0 | 0.0001 | 0 | 0 | 0 | 3e-05 |
| Red Dark Long-Lived | 0 | 0 | 0 | 0 | 0 | 0.02 | 0 | 0 | 0 | 3e-05 |

D

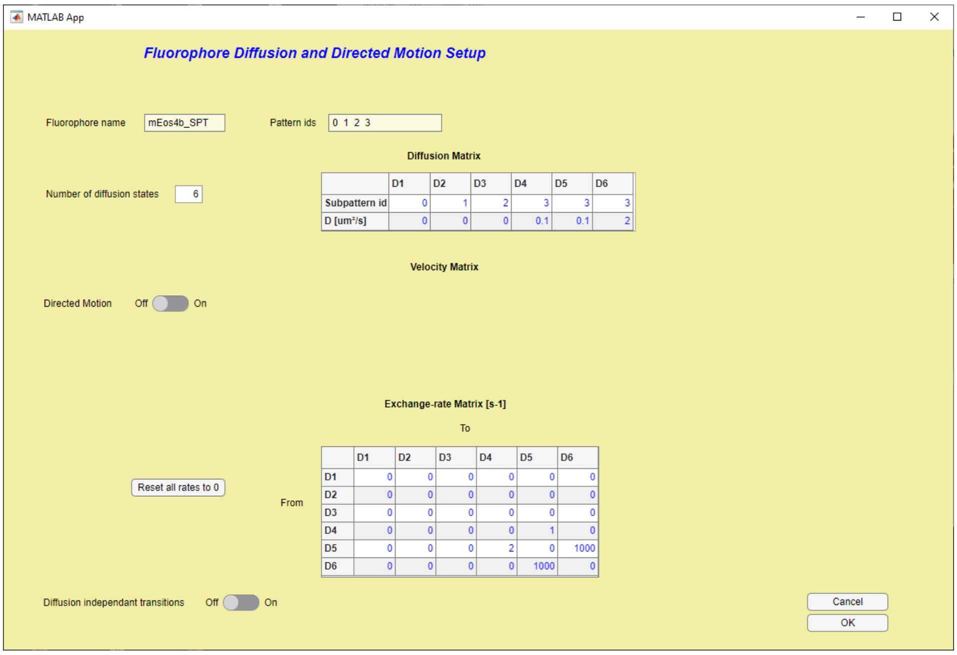


### Supplementary Fig. S26

The SMIS graphical user interface. (A) Main SMIS panel. (B) Photophysical setup panel. (C) Phototransformation quantum yields setup panel. (D) SPT setup panel.
